# Molecular characterization of the type VI secretion system effector Tlde1a reveals a structurally altered LD-transpeptidase fold

**DOI:** 10.1101/2022.08.05.502956

**Authors:** Neil Lorente Cobo, Stephanie Sibinelli-Sousa, Jacob Biboy, Waldemar Vollmer, Ethel Bayer-Santos, Gerd Prehna

**Affiliations:** Department of Microbiology, University of Manitoba, Winnipeg, MB R3T 2N2 Canada; Department of Microbiology, Biomedical Sciences Institute, University of São Paulo, São Paulo 05508-900, Brazil; Centre for Bacterial Cell Biology, Biosciences Institute, Newcastle University, Newcastle upon Tyne, NE2 4AX, United Kingdom

**Keywords:** Effector, Tlde1a, Type VI secretion system, LD-transpeptidase, LD-carboxypeptidase, peptidoglycan, *Salmonella* Typhimurium, structural biology, X-ray crystallography

## Abstract

The type VI secretion system (T6SS) is a molecular machine that Gram-negative bacteria have adapted for multiple functions, including interbacterial competition. Bacteria use the T6SS to deliver protein effectors into adjacent cells to kill rivals and establish niche dominance. Central to T6SS mediated bacterial competition is an arms race to acquire diverse effectors to attack and neutralize target cells. The peptidoglycan has a central role in bacterial cell physiology, and effectors that biochemically modify peptidoglycan structure effectively induce cell death. One such T6SS effector is Tlde1a from *Salmonella* Typhimurium. Tlde1a functions as an LD-carboxypeptidase to cleave tetrapeptide stems and as an LD-transpeptidase to exchange the terminal D-alanine of a tetrapeptide stem with a noncanonical D-amino acid. To understand how Tlde1a exhibits toxicity at the molecular level, we determined the X-ray crystal structure of Tlde1a alone and in complex with D-amino acids. Our structural data revealed that Tlde1a possesses a unique LD-transpeptidase fold consisting of a dual pocket active site with a capping subdomain. This includes an exchange pocket to bind a D-amino acid for exchange and a catalytic pocket to position the D-alanine of a tetrapeptide stem for cleavage. Toxicity assays in *Escherichia coli* and *in vitro* peptidoglycan biochemical assays with Tlde1a variants, correlate Tlde1a molecular features directly to its biochemical functions. We observe that the LD-carboxypeptidase and LD-transpeptidase activities of Tlde1a are both structural and functionally linked. Overall, our data highlights how an LD-transpeptidase fold has been structurally altered to create a toxic effector in the T6SS arms race.

## INTRODUCTION

Type six secretion systems (T6SSs) are dynamic nanomachines that Gram-negative bacteria use for numerous biological functions. These diverse functions include killing competing bacteria (1-5), self-recognition (6,7), pathogenesis (8,9), micronutrient acquisition (10-13), and direct communication by the microbiota with a eukaryotic host (14,15). Although the core proteins of the T6SS ejection apparatus are conserved across Gram-negative bacteria (16,17), the T6SSs of individual species typically harbor a unique set of protein effectors. These species unique effectors can aid in host adaptation (18,19) and are reflective of a toxin arms race between Gram-negative bacteria that compete for the same niche and resources (2,9).

At the molecular level, the T6SS forcibly secretes toxic protein effectors directly into adjacent cells in a contact dependent manner (1,3,20,21). The T6SS is structurally similar to the T4 phage tail apparatus and is organized into three main complexes: the trans-membrane, the baseplate, and the tail complex (17). The trans-membrane complex spans the periplasm anchoring the T6SS to the bacterial envelope and allows consequent docking of the baseplate (22). The baseplate serves to link the tail complex to the trans-membrane complex, while also providing an environment for effector loading onto the tip of the tail complex (17,23). The tail complex is comprised of haemolysin co-regulated protein (Hcp) hexamers enveloped in a dynamic protein sheath, which together span the bacterial cell before firing (17,24,25). Within the baseplate at the tip of the tail complex, a valine-glycine repeat protein G (VgrG) binds directly to the Hcp proteins of the tail (26). Depending upon the biological role of a particular T6SS, the individual effectors may bind directly to the VgrG spike protein or the Hcp proteins of the tail for secretion (27-29). Additionally, the mechanistic differences of T6SS effector loading can denote if the secreted toxin will localize to the cytosol or periplasm of a Gram-negative competitor (29,30).

T6SS effectors have a wide range of toxic biochemical activities. Examples include nucleases (31,32), lipases (33,34), NADases (35), ADP-ribosylstransferases (36), and various classes of peptidoglycan cleaving or modifying enzymes (5,37,38). Additionally, these effectors are often co-expressed with an immunity protein that directly binds the effector to neutralize its enzymatic function for self-protection. In many Gram-negative bacteria, adjacent cells battle each other in a ‘tit for tat’ matter with secreted volleys of their unique set of T6SS effectors (1). If the cells contain the same set of effectors and cognate immunity proteins both cells live. If their T6SS effectors differ, one species may compete the other and establish niche dominance (2,39).

A prominent example of the T6SS effectors arms race can be seen in the evolution of the numerous effectors that target the peptidoglycan (40). In Gram-negative bacteria, the peptidoglycan layer within the periplasm envelops the cytoplasmic membrane to maintain cell shape and prevent bursting due to the osmotic pressure (41,42). As such, toxin-catalyzed degradation of the peptidoglycan layer causes morphological changes to the cell that results in growth inhibition and eventual cell lysis (5,37,38).

The peptidoglycan structure consists of alternating N-acetylglucosamine (GlcNAc) and N-acetylmuramic acid (MurNAc) glycan residues that are crosslinked via short peptides. In most Gram-negative bacteria, newly synthesized peptides have the specific linkage order of L-Alanine^1^ (L-Ala), D-isoglutamic acid^2^ (D-iGlu), meso-diaminopimelic acid^3^ (mDAP), D-Alanine^4^ (D-Ala), and D-Alanine^5^ (D-Ala) (42). To promote structural stability and rigidity of the peptidoglycan later, peptides protruding from adjacent glycan chains are crosslinked by different classes of transpeptidases (TPases). DD-TPases form a 4-3 crosslink between D-Ala^4^ and mDAP^3^, whereas LD-TPases form a 3-3 crosslink between mDAPs^3^ from two tetrapeptides (43). DD-TPases, also known as penicillin binding proteins (PBPs), contain a conserved catalytic serine residue that removes D-Alanine^5^ from one peptide stem creating an acyl-enzyme intermediate. The 4-3 crosslink is formed through nucleophilic attack by an amino group of mDAP^3^ on a neighboring peptide stem regenerating the enzyme (43,44). LD-TPases catalyze a 3-3 crosslink through nucleophilic attack on one stem, which releases D-Ala^4^ and any subsequent moieties in the stem. This forms an acyl-enzyme intermediate via a conserved catalytic cysteine residue. Finally, the LD-TPase is deacylated by nucleophilic attack from the amino group of a neighboring mDAP^3^ from the acceptor tetrapeptide (43,45).

Given the complexity of the peptidoglycan structure, T6SS harboring bacteria have evolved effectors that target different molecular features of the peptidoglycan as part of the bacterial competition strategies (40). The characterized Tae (type VI amidase effector) family of effectors from *Pseudomonas* species and the Ssp enzymes from *Serratia* species act as amidases that cleave within peptides (3,4,46). Additionally, specific amidase effectors have evolved to cleave between D-iGlu^2^ and mDAP^3^ (Tae1, Ssp1) or mDAP^3^ and D-Ala^4^ within a crosslink (Tae2, Tae3) (4). Tge (type VI glycoside hydrolase effector) effectors act as glycoside hydrolases, which cleave within the glycan chains (3,47). Similar to amidase effectors, variations to the T6SS effectors denote a substrate specificity to either cleave the glycan linkage between GlcNAc and MurNAcNAM, or MurNAc and GlcNAc (40). Another T6SS effector family termed Tlde1, was recently found in several *Salmonella* serovars and in a subset of other Gram-negative bacterial species. Tlde1 (type VI LD-transpeptidase effector 1) effectors possess dual enzymatic activities, as they are able to function both as LD-carboxypeptidases (trimming of tetrapeptides to tripeptides) and LD-transpeptidases (breaking and forming new peptide bonds) (38).

Tlde1a is encoded as an effector-immunity pair with its cognate immunity protein Tldi1a within the SPI-6 (Salmonella pathogenicity island 6) T6SS in the *Salmonella* Typhimurium genome (38,48). Previous work has shown that Tlde1a has the ability to cleave muropeptides (disaccharide peptide subunits) between mDAP^3^ and D-Ala^4^ functioning as a LD-carboxypeptidase (CPase). Additionally, Tlde1a can also act as an LD-TPase, with the specialized function of exchanging the D-Ala^4^ of the tetrapeptide with another D-amino acid (38). Specifically, experiments have demonstrated that Tlde1a can degrade GlcNAc-MurNAc--tetrapeptide isolated from *E. coli* peptidoglycan into GlcNAc-MurNAc-tripeptides. Moreover, in the presence of D-methionine (D-Met) Tlde1a replaces D-Ala^4^ of GlcNAc-MurNAc-tetrapeptides with D-Met creating a new GlcNAc-MurNAc--tetrapeptide that the standard CPases of the cell-wall recycling machinery might not recognize (38,49). Overall, the dual activities of Tlde1a have the net effect hindering cell-wall homeostasis, which results in cell swelling and death.

Tlde1a contains a catalytic motif with a conserved cysteine similar to LD-TPases, <u>H</u>XX14-17-(S/T)HG<u>C</u>h (underline is conserved catalytic, X is variable, h is hydrophobic) (38). Given that Tlde1a represents a new class of T6SS effector that was recently described, we undertook a structural approach to reveal the molecular features that allow Tlde1 enzymes to possess both LD-CPase and LD-TPase exchange activities. Here we determined high resolution X-ray structures of Tlde1a from *S*. Typhimurium alone and in complex with D-Ala and D-Met. We observe that Tlde1a has a structurally altered LD-TPase domain that includes a capping subdomain. Structure guided point mutations to analyze target cell toxicity and peptidoglycan degradation revealed the molecular features of Tlde1a that provide its dual enzymatic activities. Additionally, biochemical experiments show that the LD-CPase and LD-TPase exchange activities are structurally and functionally linked. Moreover, enzymatic assays with peptidoglycan revealed that Tlde1a can also perform LD-TPase reactions with a tetrapeptide acceptor, generating 3-3 small amounts of crosslinks. Together, our results demonstrate that Tlde1a contains a single large active site cleft bounded by a capping subdomain for muropeptide binding, with two smaller pockets surrounding the catalytic cysteine. One pocket positions the terminal D-Ala^4^ of a tetrapeptide stem for cleavage, and the second pocket accepts an incoming D-amino acid for exchange and transpeptidation.

## RESULTS

### A crystal structure of Tlde1a reveals a highly conserved active site cleft

To probe the molecular details of Tlde1a, purified Tlde1a from *S*. Typhimurium was crystallized, and its structure determined using x-ray crystallography (Fig. 1 and Table 1). Although the cognate immunity protein Tldi1a was included in a co-expression construct with Tlde1a, the immunity protein failed to co-purify with the his-tagged effector. Tlde1a crystallized as a monomer in the space group P4_1_2_1_2 and the structure was modelled to a resolution of 1.65 Å. Initial phases were obtained by single anomalous diffraction (SAD) on a home source diffractometer using Tlde1a crystals soaked with sodium iodide. Due to the presence of β-mercaptoethanol in the buffer conditions, the active site residue C131 was modified to S,S-(2-hydroxymethyl)thiocysteine (CME). Complete data collection and refinement statistics are displayed in Table 1.

**Table 1.** X-ray data collection and refinement statistics.

|  | Tlde1a (native) | Tlde1a:D-Ala | Tlde1a:D-Met | Tlde1a WT NaI |
| --- | --- | --- | --- | --- |
| **Data Collection** |  |  |  |  |
| Wavelength (Å) | 1.5418 | 1.1806 | 1.5418 | 1.5418 |
| Space group | P4<sub>1</sub>2<sub>1</sub>2 | P4<sub>1</sub>2<sub>1</sub>2 | P4<sub>1</sub>2<sub>1</sub>2 | P4<sub>1</sub>2<sub>1</sub>2 |
| Cell dimensions |  |  |  |  |
| *a*, *b*, *c* (Å) | 59.56 59.56<br>106.17 | 60.14 60.14<br>105.70 | 59.46 59.46<br>106.80 | 59.60 59.60<br>103.90 |
| $\alpha$, $\beta$, $\gamma$ (°) | 90.00 90.00<br>90.00 | 90.00 90.00<br>90.00 | 90.00 90.00<br>90.00 | 90.00 90.00<br>90.00 |
| Resolution (Å) | 42.12 -1.65<br>(1.68-1.65) | 42.53 – 1.80<br>(1.84-1.80) | 42.05 – 1.60<br>(1.66-1.60) | 42.18 – 1.85<br>(1.89-1.85) |
| Total reflections | 295181 (14556) | 223732 (12626) | 341238 (16443) | 191492 (10232) |
| Unique reflections | 23788 (1129) | 18721 (1081) | 26027 (2538) | 16864 (992) |
| CC(1/2) | 0.999 (0.525) | 0.993 (0.343) | 0.999 (0.961) | 0.997 (0.546) |
| *R*<sub>merge</sub> | 0.077 (1.829) | 0.185 (1.693) | 0.062 (0.449) | 0.069 (1.760) |
| *R*<sub>pim</sub> | 0.032 (0.752) | 0.078 (0.737) | 0.025 (0.181) | 0.030 (0.804) |
| *I*/σ*I* | 16.1 (1.0) | 9.1 (1.2) | 23.9 (4.4) | 24.1 (1.4) |
| Completeness (%) | 100.0 (100.0) | 100.0 (100.0) | 99.25 (99.80) | 99.8 (98.4) |
| Redundancy | 12.4 (12.9) | 12.0 (11.7) | 13.1 (13.2) | 8.4 (10.3) |
| **Refinement** |  |  |  |  |
| *R*<sub>work</sub> / *R*<sub>free</sub> (%) | 16.34 / 18.82 | 18.82 / 21.53 | 18.90 / 21.06 |  |
| Average B-factors (Å<sup>2</sup>) | 27 | 41.21 | 25.59 |  |
| Protein | 25.25 | 40.34 | 24.08 |  |
| Ligands | 56.12 | 62.8 | 25.84 |  |
| Water | 34.36 | 44.3 | 33.91 |  |
| No. atoms | 1677 | 1619 | 1709 |  |
| Protein | 177 | 178 | 180 |  |
| Ligands | 25 | 44 | 24 |  |
| Water | 34.36 | 44.3 | 33.91 |  |
| Rms deviations |  |  |  |  |
| Bond lengths (Å) | 0.009 | 0.004 | 0.011 |  |
| Bond angles (°) | 0.95 | 0.64 | 1.02 |  |
| Ramachandran plot (%) |  |  |  |  |
| Total favored | 98.24 | 98.25 | 98.84 |  |
| Total allowed | 1.76 | 1.75 | 1.16 |  |
| PDB code | 7UMA | 7UO3 | 7UO8 |  |

Statistics for the highest-resolution shell are shown in parentheses

**Figure 1:**
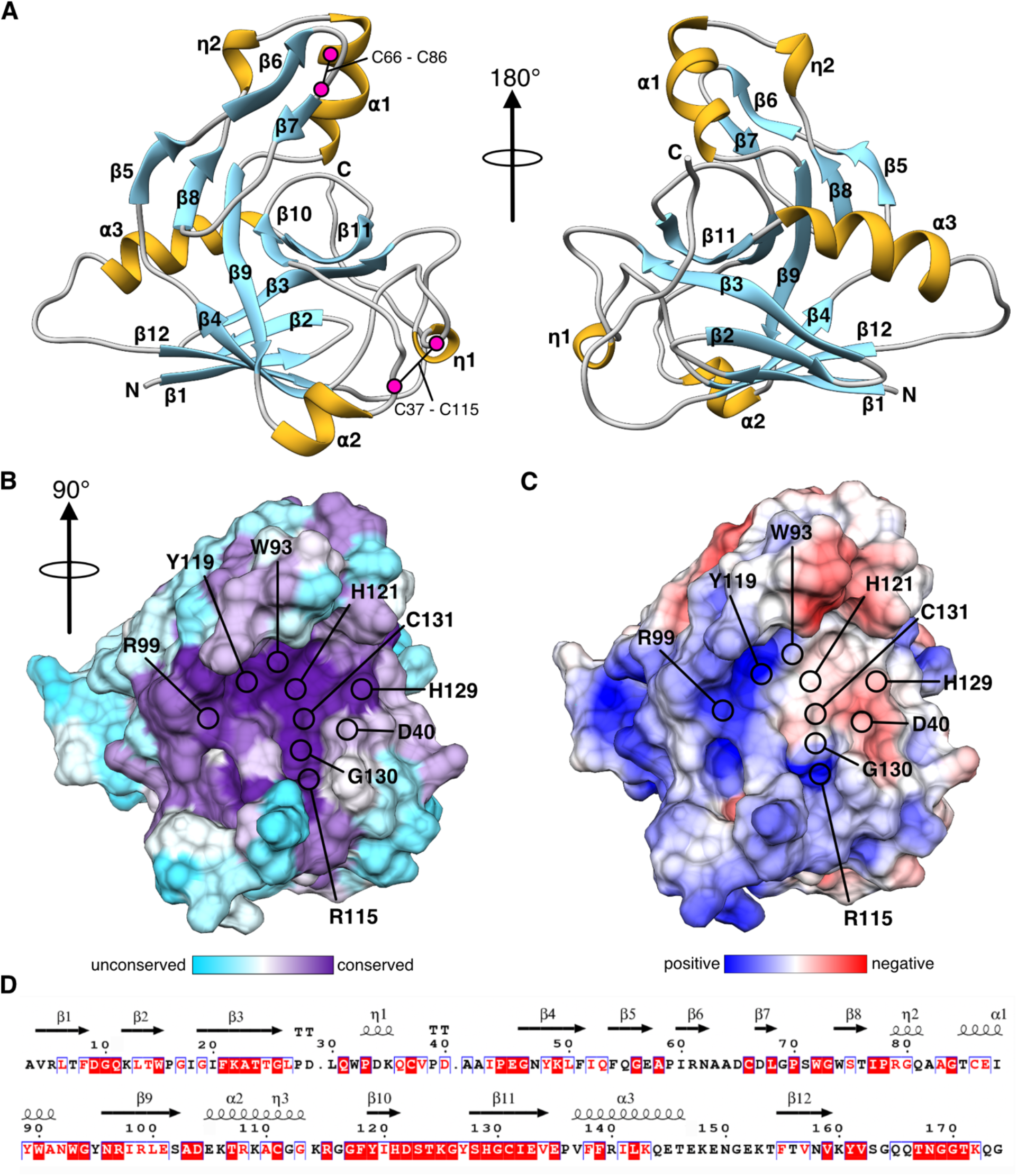
X-ray crystal structure of Tlde1a. A) Two views of a cartoon ribbon depiction of *S*. Typhimurium Tlde1a. Secondary structure is labeled by alpha-helix (α), 3_10_ helix (η), and beta-strand (β) with the N-terminus and C-terminus labeled. Positions of conserved disulphide bonds are indicated by residue number and magenta circles. B) Active site surface of Tlde1a colored by residue conservation. Conserved residues in the putative substrate binding pocket or that comprise the catalytic motif are highlighted C) Active site surface of Tlde1a colored by electrostatic potential. Electrostatic potential was calculated at pH 7.0 and is scaled between -5 kT/e to 5 kT/e. The same residues are highlighted as in B. D) Conserved residues of Tlde1a plotted with secondary structure. Red highlights are strict conservation, red letters indicate homologous residues, and black letters are unconserved. Multisequence alignment was generated with Consurf (https://consurf.tau.ac.il/) and plotted by Espript (https://espript.ibcp.fr/).

As shown in Figure 1A, Tlde1a adopts a primarily β-sheet fold (blue) with the core of the domain comprised of a β-sandwich (β1-4 and β9-12) that is bounded on either side by α-helices (α2 and α3) (gold). The side of the β-sandwich opposite of helix α2 is also part of a continuous β-sheet (β5, β8-11, β3) that faces a small sub-domain consisting of strands β6-7, helix α1, and the second 3_10_ helix (η2). Additionally, the Tlde1a structure is stabilized by two conserved disulfide linkages highlighted in magenta (C66-C86 and C37-C115). Furthermore, each disulfide bond links amino acids that are 20 and 78 positions apart in the Tlde1a primary sequence. Of particular note is that the conserved LD-TPase residue motif (<u>H</u>XX14-17-(S/T)HG<u>C</u>h) is part of the continuous β-sheet structure (β10-11) and connecting loop regions (Fig. 1A). This creates an active site cleft bounded by the β6-7 insertion sub-domain and the loop region containing helix α2 and the first 3_10_ helix (η1).

A surface representation of Tlde1a colored by residue conservation and electrostatic potential is shown in Figure 1B and Figure 1C, respectively. Residue conservation was plotted based on results of the Consurf server (50) and electrostatics calculated using the adaptive Poisson-Boltzmann solver (51). Each of these views is rotated 90° relative to Figure 1A to look directly into the active site. As clearly demonstrated, Tlde1a folds to create a significant active site cleft that is highly conserved. This includes H121, H129, and C131 that are part of the LD-TPase residue motif (<u>H</u>XX14-17-(S/T)HG<u>C</u>h) (38) and a number of other conserved residues whose positions have been indicated on the Tlde1a structure. These residues include polar residues (R99, R115), which could serve as hydrogen bond donors, and aromatic residues (W93 and Y119) that may provide van der Waals packing with D-Ala sidechains of a peptidoglycan tetrapeptide. The displayed electrostatic characteristics of Tlde1a indicate that the right side of the active site cleft near residue C131 carries negative potential, whereas the outer side of the cleft near residue R99 trends towards positive (Fig. 1C). This observation supports the observed LD-CPase and LD-TPase activities, where the C131 thiol must be deprotonated in order to catalyze cleavage of a tetrapeptide. Furthermore, a hydrophobic surface analysis of Tlde1a is shown in Figure S1A, which suggests that the opposite electrostatic potentials of the active site cleft are separated by a hydrophobic patch. Additionally, the partial hydrophobic nature of the Tlde1a active site may serve to bind D-Ala^4^ residue sidechains of a tetrapeptide stem. Figure 1D plots the residue conservation of Tlde1a relative to secondary structure elements, with an alignment of the top five homologues from the Consurf multisequence alignment shown in Figure S1B.

### Tlde1a adopts a structurally unique LD-transpeptidase fold

When analyzed by the Dali server (52), Tlde1a shows high homology to the catalytic domain of several LD-TPase enzymes such as *Mycobacterium tuberculosis* LdtMt2 (PDBid 4GSQ) (53) and *E. coli* LdtD (YcbB, PDBid 6NTW) (54), in addition to the *Helicobacter pylori* LD-CPase Csd6 (PDBid 4XZZ) (55). Alignment of Tlde1a with these structural homologues shows that the core β-sandwich and extended β-sheet along with the α2-helix of Tlde1a make up the characteristic elements of the LD-TPase fold (Fig. 2). However, unlike Tlde1a other peptidase enzymes appear covalently linked to additional functional domains. For example, LdtMt2 has an N-terminal domain reminiscent of an Ig-fold (53), YcbB possess a PG-domain for peptidoglycan binding (54), and Csd6 has an NTF-like domain that may have pseudoaminidase activity (55) (Fig. 2). Furthermore, in contrast to enzymes such as YcbB, Tlde1a does not require a secondary PG-domain for its enzymatic function.

**Figure 2:**
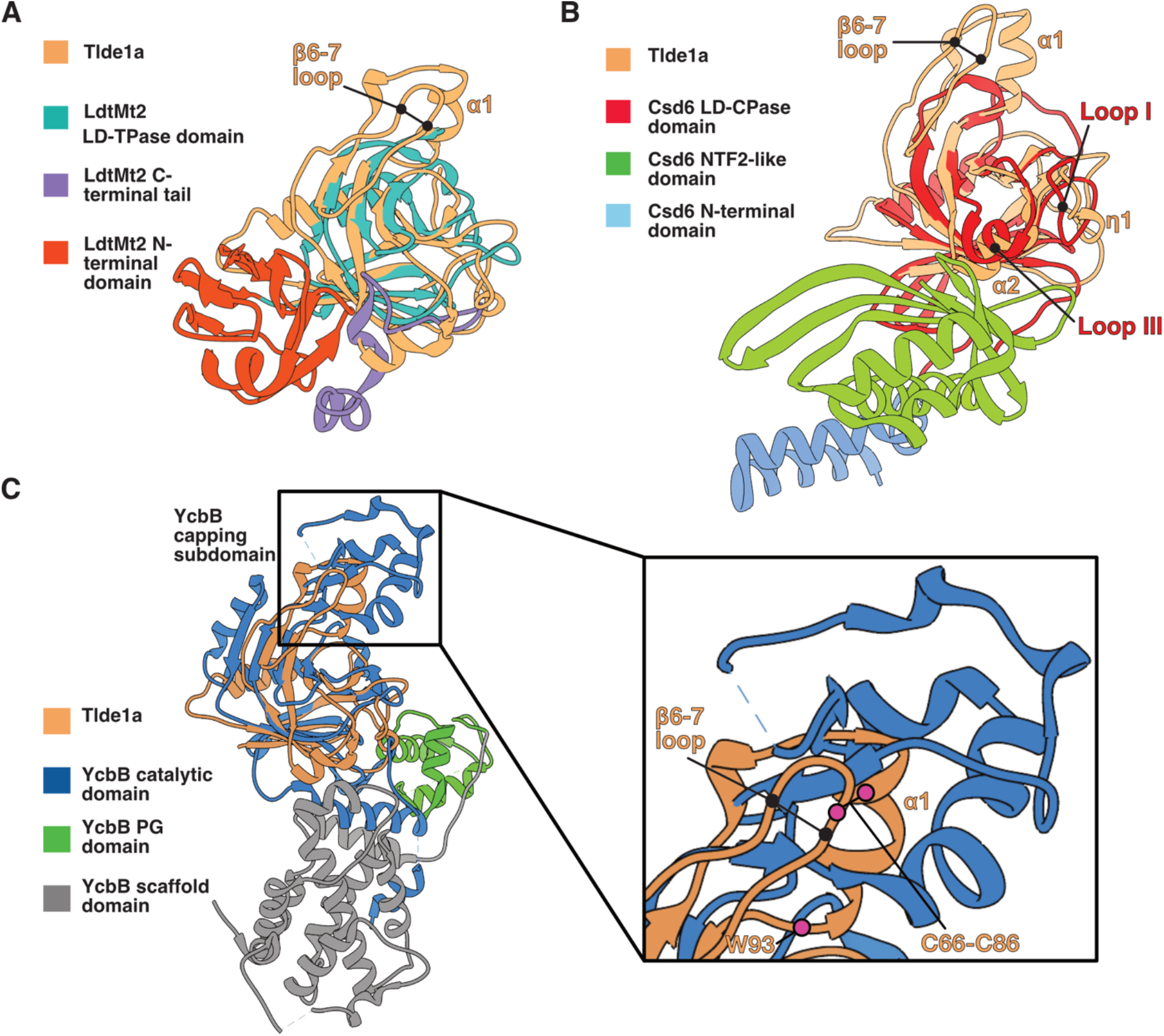
Comparison of Tlde1a with structural and functional homologues. A) Overlay of Tlde1a with the LD-transpeptidase domain of LdtMt2 (PDBid 4GSQ). B) Overlay of Tlde1a with the catalytic domain of the LD-carboxypeptidase Csd6 (PDBid 4XZZ). Csd6 loops I and III are indicated in relation to η1 and α2 of Tlde1a, respectively. C) Overlay of Tlde1a with the catalytic domain of the LD-transpeptidase YcbB (PDBid 6NTW). Alignment of the Tlde1a capping subdomain with the YcbB capping subdomain is highlighted. The insert displays a closer view of the capping domains showing core secondary structure elements and conserved residues of Tlde1a. For each panel Tlde1a is colored in beige and each structural homologue is colored by functional domain. Each panel also indicates the positions of the β6-7 loop and α-helix that comprise the Tlde1a capping subdomain.

Comparing the core LD-TPase fold of Tlde1a to structural homologues we observe that Tlde1a has a unique fold. Relative to LdtMt2 and Csd6, Tlde1a contains an extra small subdomain (β6-7, helix α1, and 3_10_ helix (η2)) (Fig. 1A, Fig. 2A and Fig. 2B). Similarly, YcbB also has a subdomain known as the ‘capping subdomain’ in the same structural position as Tlde1a relative to other LD-TPase enzymes (Fig. 2C). However, the YcbB capping subdomain is significantly larger than the Tlde1a subdomain and displays a different protein fold. The YcbB capping subdomain has been shown to form a cap over its substrate binding cleft and has been hypothesized to assist in binding peptidoglycan substrates (54). Interestingly, the Tlde1a subdomain appears to provide a similar structural role relative to the Tlde1a active site (Fig. 1A-B).

As Tlde1a has dual enzyme activities, we compared the active site cleft of Tlde1a relative to a structurally similar LD-CPases. For this analysis we primarily focused on Csd6, as Csd6 it is the closest structural homolog of Tlde1a as determined by the Dali server (Z-score 11). Additionally, Csd6 has a canonical LD-TPase domain that only exhibits LD-CPase activity. Structural comparison of Csd6 to LD-TPases domains highlighted that loop I and loop III differed from the canonical LD-TPases fold and that they could be a determinant of the LD-CPase activity of Csd6 (55). As shown in Figure 2B, Tlde1a appears to also have similar molecular features, represented by the α2 and the 3_10_ helix (η1). As this region of Tlde1a is locally stabilized by a conserved disulfide bond (C37-C115) (Fig. 1A), we hypothesize that the α2 and the 3_10_ helix (η1) loop region may contribute to the enzymatic activity of Tlde1a.

### The active site cleft of Tlde1a contains a binding pocket for D-amino acids

Past work has demonstrated that Tlde1a uses its LD-TPase activity to exchange the terminal D-Ala^4^ of a donor tetrapeptide stem with D-amino acids that are not normally part of newly synthesized peptidoglycan (38). To probe the molecular features of Tlde1a that may contribute to this activity, we soaked Tlde1a crystals with D-Ala and D-Met. We were able to obtain co-crystal structures of Tlde1a in complex with each amino acid. Additionally, all structures also include a crystal packing artifact of the Tlde1a C-terminal His-tag from a symmetry mate within the active site (Fig. 3 and Table 1).

**Figure 3:**
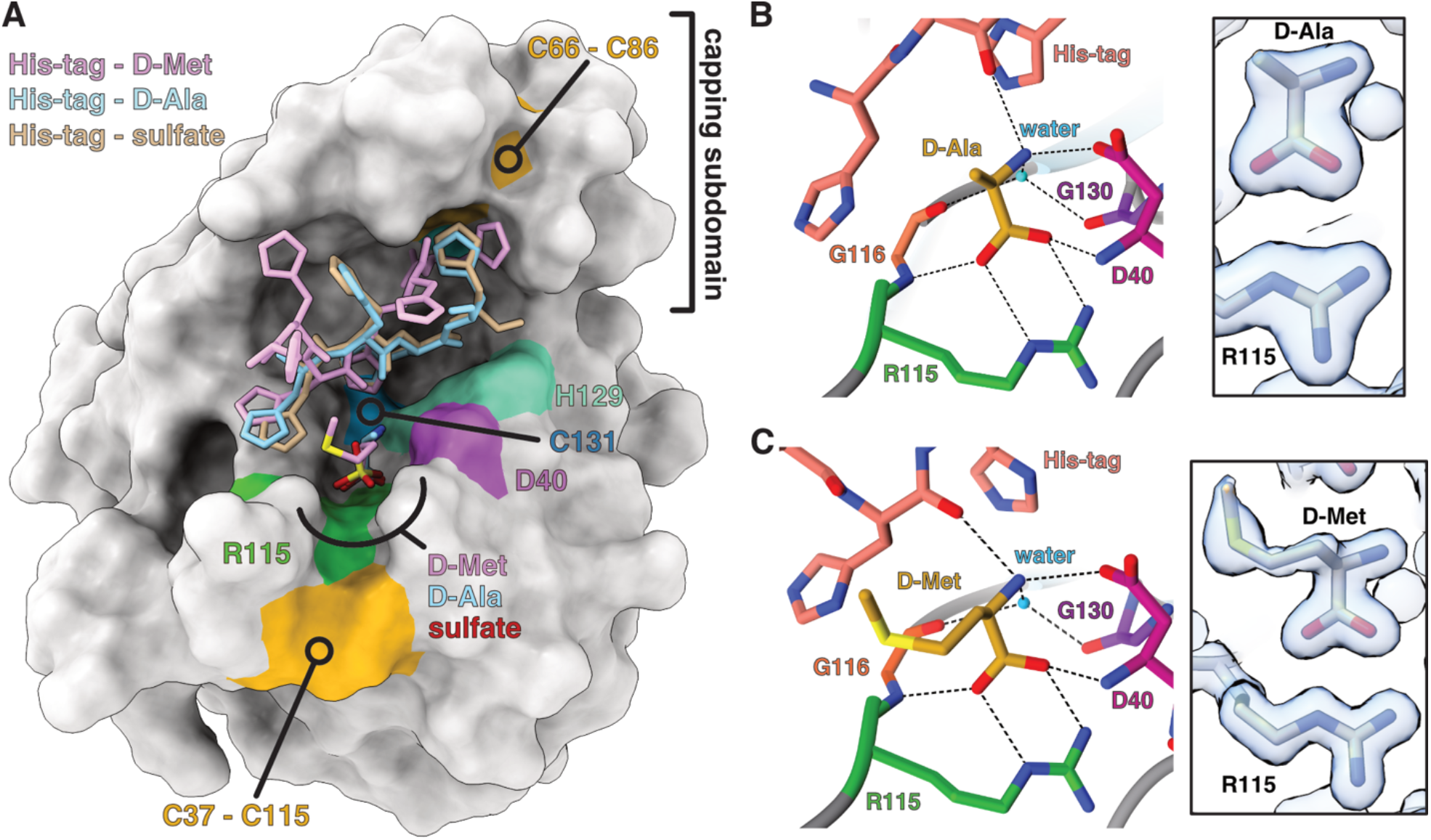
X-ray co-crystal structures of Tlde1a bound to D-alanine and D-methionine. A) Surface representation of Tlde1a with ligands overlaid. The positions of bound sulfate (SO_4_^2-^) (PDBid 7UMA), D-Alanine (D-Ala) (PDBid 7UO3), and D-Methionine (D-Met) (PDBid 7UO8) in addition to the corresponding His-tag cloning artifact captured in the active site are shown and labeled by color. The positions of conserved active site residues are also shown and labeled by color. B) Molecular details of free D-Ala binding and C) free D-Met binding by Tlde1a. Residues that interact with each D-amino acid are labeled by color and number, with hydrogen bonds indicated by dashed lines. The observed electron density of each D-amino acid is shown contoured at 1.5 rmsd (right).

As shown in Figure 3A, Figure 3B and Figure 3C, both D-amino acids bind in the same pocket and are stabilized by an equivalent network of hydrogen bonds. The interaction of both D-amino acids with Tlde1a is facilitated by a salt-bridge between the D-amino acid carboxylic acid and the sidechain of R115. Additional H-bonding to the carboxylic acid group is provided by the backbone amides of D40 and G116. Finally, the amide group of the D-amino acid participates in a bifurcated hydrogen bond with the acidic sidechain of D40, a water molecule, and a carbonyl of the Tlde1a His-tag cloning artifact. This ordered water is also H-bonded to the backbone carbonyls of G116 and G130. Furthermore, the observed water is positioned ∼3.7 Å from C131 and is the closest ordered solvent molecule to C131 in the active site. Interestingly, the hydrophobic sidechain of D-Met faces towards the solvent and has no significant interaction with the active site of Tlde1a. Moreover, in the Tlde1a native structure the D-amino acid binding pocket is occupied by a sulfate from the crystallization condition supporting the observation that H-bonds are the primary mode of ligand binding (Fig. 3A and Fig. S2A). It is also important to note that the Tlde1a complex with D-Ala contains a second bound D-Ala. However, this D-Ala is at a crystal packing interface, interacts with unconserved residues, and shows a poorer fit to the electron density (Fig. S2B). Taken together, these observations suggest that Tlde1a contains a D-amino acid binding pocket that may have no sidechain selectivity for the D-amino acid to be exchanged.

### A crystal packing artifact provides a model for peptide binding

All three Tlde1a crystal structures included the His-tag of a symmetry mate bound in the Tlde1a active site as a crystal packing artifact (Fig. 3 and Fig. S2C-E). For the native and D-Ala bound structures four residues of the 6His-tag could be modeled whereas for the D-Met co-crystal complex five histidines were observed in the electron density. In all three structures the His-tag occupies the highly conserved active site and makes van der Waals packing contacts with the Tlde1a capping subdomain (β6-7, helix α1, and 3_10_ helix (η2))). However, most of the peptide backbone interactions of the His-tag with Tlde1a are mediated by ordered water molecules or sulfates from the crystallization condition (Fig. S2A). Although the His-tag is an artifact, binding of the known D-Met substrate to Tlde1a induces a drastic conformational change in the His-tag. Compared to the native and D-Ala bound structures, the His-tag in the D-Met bound structure has significantly shifted its position within the Tlde1a active site (Fig. 3A). This change also includes the ability to model 5 His-tag residues instead of 4 (Fig. S2C-E) and the loss of two ordered sulfates (Fig. S2A). Additionally, the binding of D-Met and the re-ordering of the His-tag induces a conformational change in the Tlde1a capping subdomain relative to both the sulfate and D-Ala bound structures (Fig. S2F). As the capping subdomain of YcbB undergoes conformational changes to accommodate its peptidoglycan substrates (54), we hypothesize that the His-tag artifact approximates how Tlde1a initially recognizes a peptide.

### The Tlde1a active site contains two distinct pockets

To further model how Tlde1a binds its substrates, we attempted to dock peptidoglycan tetrapeptides in the Tlde1a active site using the program SwissDock (56,57). The modified active site CME131 was mutated to C131 for ligand modeling and the His-tag artifact removed. Docking attempts were made using two different tetrapeptides (pubchem CIDs 25201603 and 134820170) and the results of the simulations are shown in Figure 4, Figure S3, and Figure S4. Pubchem CID 25201603 is the Gram-negative pentapeptide L-Ala^1^-γGlu^2^-mDAP^3^-D-Ala^4^-D-Ala^5^ that was modified to L-Ala^1^-γGlu^2^-mDAP^3^-D-Ala^4^, and pubchem CID 25201603 is the Gram-positive tetrapeptide L-Ala^1^-γGlu^2^-D-Lys^3^-D-Ala^4^. Our *in silico* experiments yielded several binding configurations for both tetrapeptides within the active site of Tlde1a of roughly equivalent quality (Fig. S3A and B, Fig. S4A and B). Furthermore, the representative clusters (cluster 8 and cluster 1) shown in Figure 4A and Figure 4B occupy a similar space in the Tlde1a active site as the His-tag artifact (Fig. S3C and Fig. S4C). For this reason, cluster 8 (Gram-negative tetrapeptide) and cluster 1 (Gram-positive tetrapeptide) were chosen as the best fit for predictive structural analysis.

**Figure 4:**
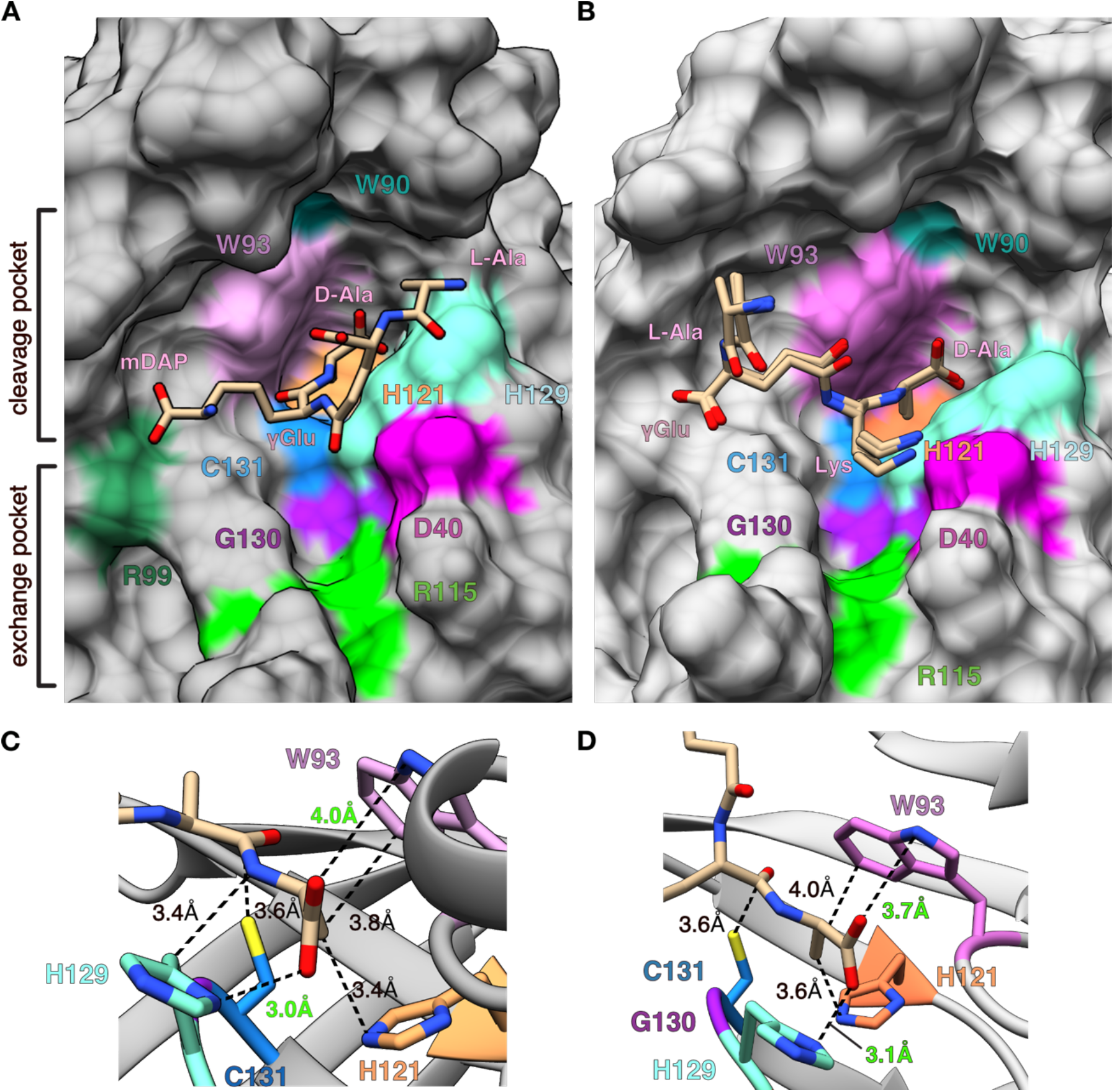
Molecular docking of peptidoglycan tetrapeptides with Tlde1a. A) Predicted binding of the tetrapeptide L-Ala^1^-γGlu^2^-mDAP^3^-D-Ala^4^ by the program Swiss-dock. An overlay of 8 similarly docked structures are shown (cluster 8). Two putative amino acid binding pockets are indicated. An ‘exchange pocket’ for the binding of a D-amino acid, and a ‘cleavage pocket’ formed by the conserved catalytic residues. B) As panel A but with the tetrapeptide L-Ala-*γ*Glu-Lys-D-Ala and cluster 1. C) Interaction of conserved Tlde1a active site residues with the docked tetrapeptide L-Ala^1^-γGlu^2^-mDAP^3^-D-Ala^4^ at the terminal D-Ala^4^. D) Interaction of conserved Tlde1a active site residues with the docked tetrapeptide L-Ala^1^-γGlu^2^-Lys^3^-D-Ala^4^ at the terminal D-Ala^4^. Distances are shown by dotted lines and labeled. van der Waal interaction distances are colored black and possible hydrogen bond interactions are colored in green.

The docked tetrapeptide ligands make several contacts with conserved residues in the Tlde1a active site. Importantly, the C-terminal D-Ala^4^ residue of each docked ligand is predicted to bind in a pocket formed by conserved LD-TPase motif catalytic residues H121, H129 and C131 (Fig. 4C and Fig. 4D). This pocket is different from the one in which D-Ala and D-Met are observed to bind in the crystal structures (Fig. 3 and Fig. 4). Additionally, the mDAP sidechain is predicted to make a hydrogen bond with R99 that appears to help position the tetrapeptide within the active site (Fig. 4A). As shown in Figure 4C and Figure 4D, the D-Ala^4^ residues of both tetrapeptides are predicted to bind deep within the pocket created by the conserved residues W93, H121, H129 and C131. In each model, the D-Ala^4^ is held in place by several van der Waals packing interactions between its aliphatic sidechain and residues W93, H121, and H129. Furthermore, the carboxylic acid group of D-Ala^4^ is positioned to make possible H-bonds with the sidechains of both W93 and H129. While the predicted H-bond with H129 is within observed hydrogen bonding distance, the predicted interaction with W93 would require a minor conformational change in W93. In total, these interactions are predicted to hold D-Ala^4^ in place such that its amide bond with the peptide stem is in position to be cleaved by C131 (see C131 distance to D-Ala in Fig. 4C and Fig. 4D). It is important to note that despite the similarities of how D-Ala^4^ is docked in each structure, the tetrapeptides are bound in different conformations. The two tetrapeptides are predicted to bind with the N-terminal D-Ala^1^ residue on opposite sides of the Tlde1a active site cleft. Based on our crystal structures and docking experiments, C131 appears to divide part of the Tlde1a active site into two distinct pockets. An ‘exchange pocket’ for binding a free D-amino acid to be swapped with the peptide stem D-Ala^4^, and a ‘catalytic pocket’ for positioning and cleavage of the terminal D-Ala^4^ for cleavage (Fig. 4A).

### The D-amino acid exchange pocket and the capping subdomain are important for Tlde1a toxicity

Based on our structural analysis, we observed molecular features of Tlde1a that may contribute to its dual enzymatic activities. Specifically, our crystal structures revealed that the Tlde1a active site contains a D-amino acid binding pocket for exchange and that the enzyme has a capping subdomain that caps the active site cleft (Fig. 1, Fig. 2, and Fig. 3). Furthermore, the His-tag artifact combined with the docking of a tetrapeptide predict the importance of a second catalytic pocket in the active site of Tlde1a (Fig. 4). Together, these results point to a number of conserved Tlde1a active site residues that likely contribute to its activity in addition to the canonical LD-TPase motif. All residues of predicted functional importance are shown on the structure of Tlde1a in Figure 1C and Figure 4A. Residues W90 and S128 were also added to our list of residues of interest due to their conservation and structural importance. The sidechain of W90 is buried within the Tlde1a capping subdomain and appears critical for proper folding. S128 is part of the LD-TPase motif and forms H-bonds with both the sidechain of H121 and the backbone amide of C131. Residue S128 was not annotated in the surface representations shown in Figure 1 as it is buried within the Tlde1a protein fold.

To test the functional importance of our structural predictions, we generated point mutation variants of Tlde1a at these residues using the plasmid pBRA (58) which contains a signal peptide (SP) *pelB* sequence for periplasm targeting (pBRA SP-Tlde1a). Alanine was chosen as the substitution for all variants except Tlde1a_G130Q_. As G130 forms the bottom of the D-amino acid exchange pocket (Fig. 4), it was replaced by Q to potentially fill the pocket. We used these plasmids to perform *E. coli* toxicity assays (Fig. 5). The results revealed that all single point mutations affect Tlde1a toxicity, except for the Tlde1_D40A_ mutation (Fig. 5A). As a control, we expressed the native version of Tlde1 (Tlde1_WT_) and Tlde1 with a point mutation in the catalytic cysteine residue (Tlde1_C131A_), which was already reported to lose its toxicity (Figure 5A) (38).

**Figure 5:**
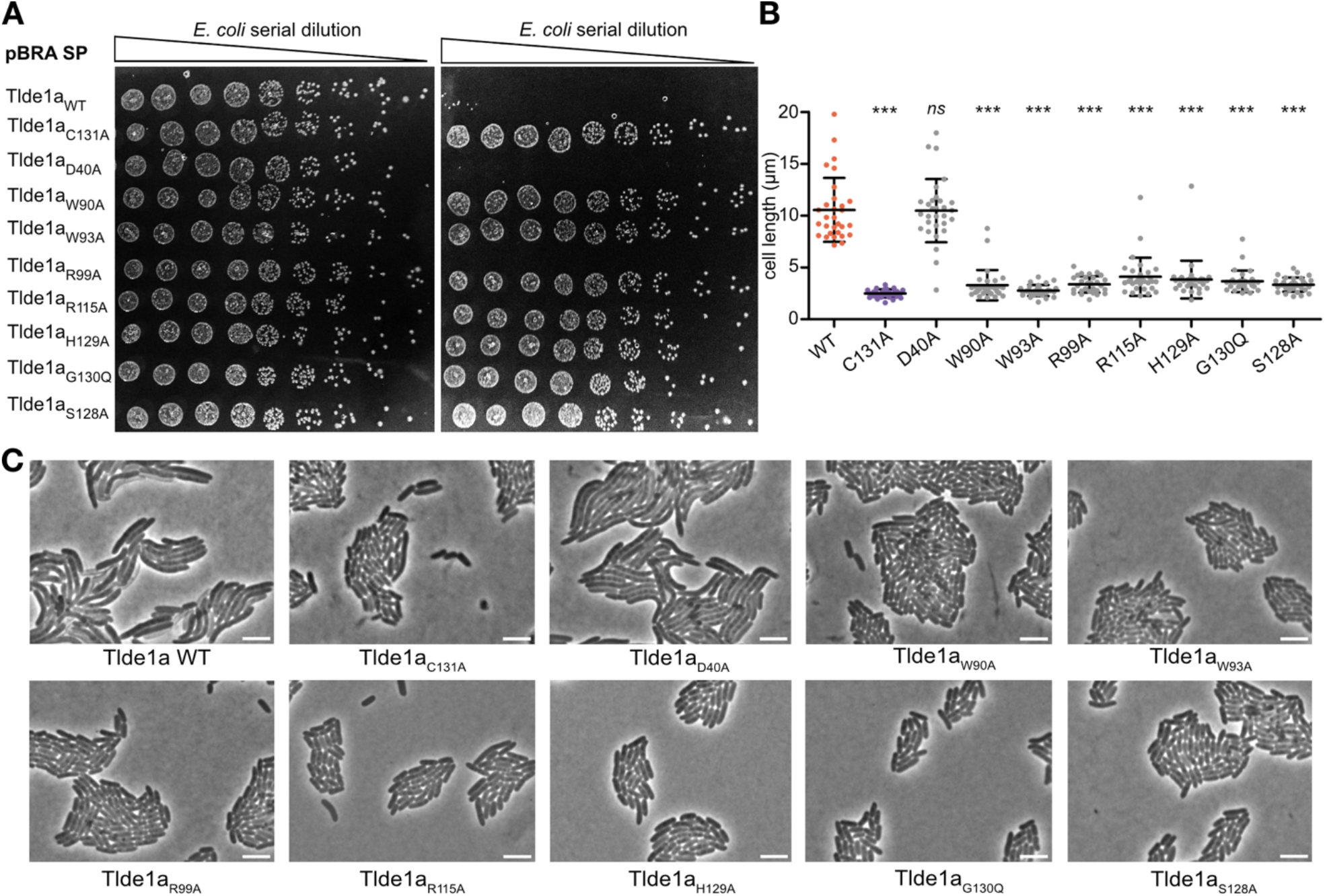
Point mutations of conserved Tlde1a residues abrogates toxicity. A) Serial dilutions of *E. coli* carrying pBRA SP-Tlde1a_WT_ or mutants spotted onto LB-agar plates containing either 0.2% D-glucose (repressed) or 0.2% L-arabinose (induced). Growth inhibition is observed upon expression of the plasmid pBRA SP-Tlde1a_WT_. Except for the mutant Tlde1a_D40A,_ all the point mutations abolished Tlde1 toxicity. The catalytic mutant Tlde1a_C131A_ was used as a control. Images are representative of three independent experiments. B) Time-lapse microscopy of *E. coli* cells carrying pBRA SP-Tlde1a_WT_ or mutants grown on LB-agarose pads containing 0.2% L-arabinose (induced). Representative images of the time point 4.5 h post-induction. Scale bar, 5 μm. C) Cell length of cells observed in B; error bars represent the SDs of the means of 30 cells measured at the time point 4.5 h post-induction. Cells carrying pBRA SP-Tlde1a mutants were analyzed through comparison with Tlde1a_WT_ by 1-way ANOVA followed by Dunnett’s multiple comparison test. *** p < 0.0001 and ns, not significant

Previously, it was observed that bacteria expressing periplasmic directed Tlde1a (pBRA SP-Tlde1a_WT_) stopped dividing or divided with a longer time interval and have a larger cell size than non-intoxicated cells (38). Moreover, the cells tend to swell and lyse, indicating that Tlde1a acts directly on the peptidoglycan (38). To verify whether this occurred with bacteria expressing Tlde1a with point mutation variants, we analyzed the growth of each strain by time-lapse microscopy. Bacteria expressing Tlde1_D40A_ tend to have a larger size, swell and lyse similarly to bacteria expressing Tlde1_WT_; however, bacteria expressing Tlde1a with the other point mutations show normal cell morphology and division (Fig. 5B and Fig. 5C) confirming their importance to Tlde1a toxicity (Videos S1-S10).

To test if the loss of function of our Tlde1a variants was not simply due to destabilization of the protein fold, we purified each point mutant and assayed their behavior in solution relative to Tlde1a_WT_. The Tlde1a gene was cloned into pET28b (38) for recombinant expression in *E. coli* and mutations were introduced. Each Tlde1a variant was readily purifiable by affinity chromatography followed by size exclusion chromatography (SEC) (Fig. 6A). However, Tlde1a_S128A_ began to degrade immediately after purification. We next monitored the wild-type enzyme and each variant by dynamic light scattering (DLS) (Fig. 6B). DLS measures an apparent particle size in solution (hydrodynamic radius) and can readily indicate protein misfolding and aggregation relative to a control sample. Most variants displayed primarily the same scattering profile in solution as Tlde1a_WT_ and closely matched its hydrodynamic radius (∼3.0 nm), except for Tlde1a_S128A_ and Tlde1a_R99A_. These results suggest that these two mutations show a loss of toxic activity due to protein misfolding and not a specific biochemical function. We conclude that the remaining highlighted residues participate in the LD-CPase and/or LD-TPase activities of Tlde1a. From this data we also conclude that the D-amino acid exchange pocket (R115) and the capping subdomain (W90) are molecular features of Tlde1a that contribute to its toxicity and thus its ability to modify the peptidoglycan.

**Figure 6:**
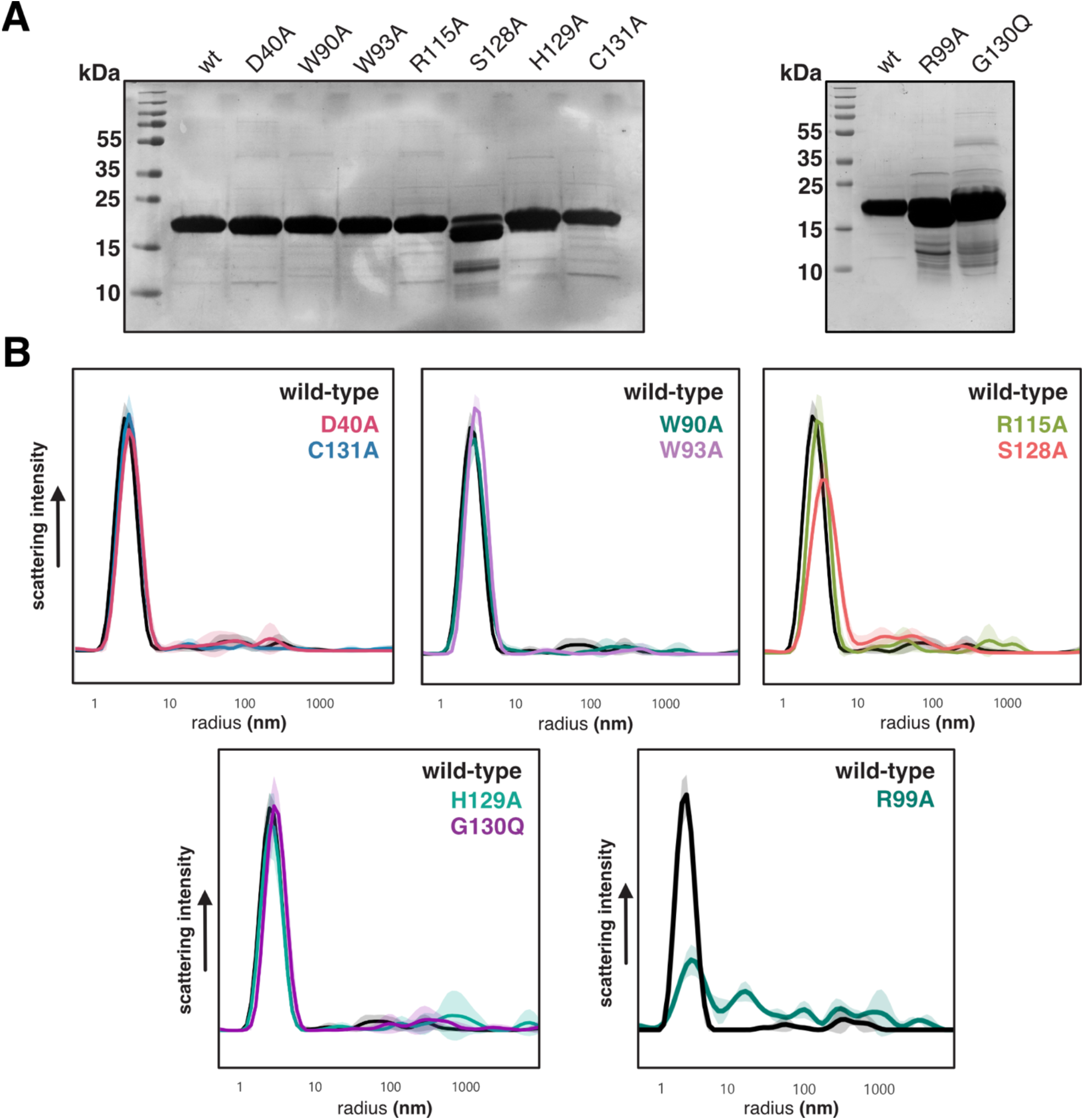
Purification and stability analysis of Tlde1a variants. A) Coomassie dye stained SDS-PAGE gels of purified Tlde1a_WT_ and Tlde1a variants used in this study. B) Dynamic light scattering of purified Tlde1a variants compared to wild-type Tlde1a. Plots represent the average of triplicate experiments. Each Tlde1a variant is shown by a colored plot of scattering intensity vs radius, with error values represented by the lighter color shade.

### LD-CPase and LD-TPase activities of Tlde1a variants against peptidoglycan

As we prepared a set of Tlde1a variants that abrogate activity and represent different molecular features of the active site, we asked how these variants might affect the ability of the enzyme to modify peptidoglycan. Specifically, we asked which variants are important for the LD-CPase activity of Tlde1a and which are important the LD-TPase exchange activity. For these assays we selected Tlde1a_D40A_, Tlde1a_R115A_ and Tlde1a_G130Q_ to represent the exchange pocket, and Tlde1a_W93A_ and Tlde1a_H129A_ to represent the catalytic pocket (Fig. 4). Tlde1a_C113A_, was included as a negative control.

For our biochemical LD-CPase and LD-TPase assays, we incubated Tlde1a_WT_ and variants with peptidoglycan from *E. coli* BW25113Δ6LDT which lacks all known proteins with a YkuD domain (LD-TPases). When peptidoglycan from *E. coli* BW25113Δ6LDT is digested into muropeptides by the enzyme cellosyl, it shows a peptidoglycan elution profile by HPLC that is dominated by the monomeric disaccharide tetrapeptide (the muropeptide Tetra, peak 3) and bis-disaccharide tetratetrapeptide (TetraTetra, peak 8) (Fig. 7A and Fig. S5). This characteristic elution profile serving as a baseline relative to Tlde1a activity. Additionally, to assay D-amino acid exchange by Tlde1a, we also incubated the peptidoglycan in the presence or absence of D-leucine (D-Leu). The amino acid D-Met was not used in these assays as some of the resulting D-Met containing muropeptides co-eluted with unmodified muropeptides, preventing correct quantitative analysis by HPLC. However, the D-Leu containing muropeptides shifted away from the unmodified muropeptides to higher retention times. Control reactions included no Tlde1a enzyme or catalytic inactive Tlde1a_C131A_ (Fig. 7A). For all assays, the muropeptides released by cellosyl were reduced with sodium borohydride and separated by HPLC (59) (Fig. 7, Fig. S5). Selected muropeptides were collected and analyzed by LC-MS/MS (Table S1).

**Figure 7:**
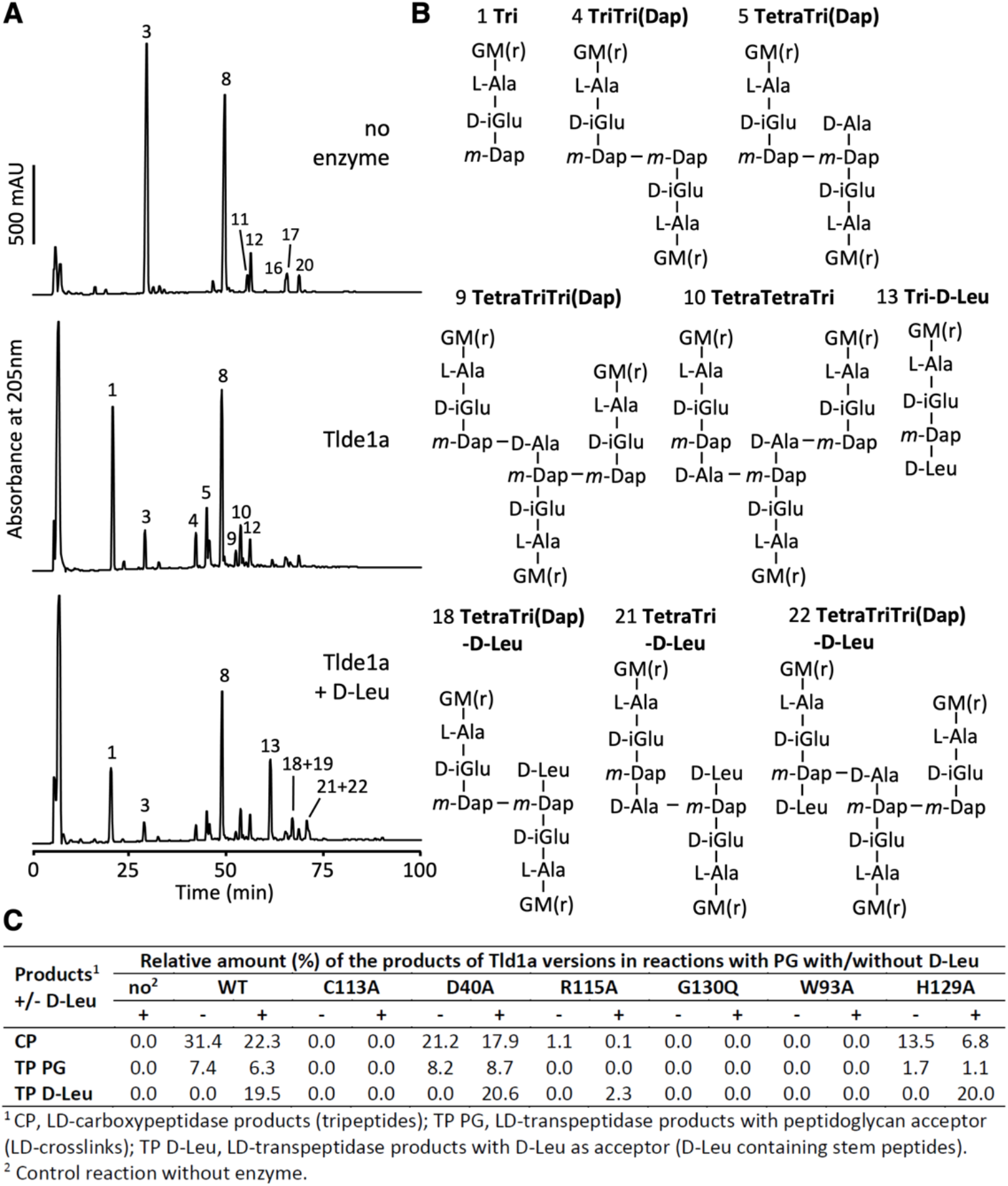
Tlde1a activity assays. A) HPLC chromatograms showing the muropeptides released from peptidoglycan after reactions of wild-type Tlde1a in the presence or absence of D-Leu and reduced with sodium borohydride. A) control reaction did not contain enzyme. The major peaks are numbered. B) Structures of the major muropeptides separated in A. Modified peaks were verified by mass spectrometry analysis (Table S1). Structures of the main unmodified muropeptides are shown in Suppl. Figure S4B. G, GlcNAc; M(r), N-acetylmuramitol; L-Ala, L-alanine; D-iGlu, D-isoglutamate; m-Dap; meso-diaminopimelic acid; D-Ala, D-alanine. C) Molar percentages of the CPase products (CP), products from LD-TPase reactions with peptide acceptors (TP PG) and products from LD-TPase reactions with D-Leu acceptor (TP D-Leu) obtained with different Tlde1a variants.

Wild-type Tlde1a was able to modify peptidoglycan by two reactions. We confirmed previous results with muropeptides showing the LD-CPase activity that produces tripeptides (e.g., muropeptide 1, Fig. 7B). In addition, we also observed the formation of 3-3 crosslinked muropeptides (muropeptide 5, Fig. 7B) which are generated by LD-TPase reactions with a peptide as acceptor. We also noticed the generation of muropeptides that must have been produced by both LD-CPase and LD-TPase reactions, e.g. muropeptide 4 (Fig. 7B). When Tlde1a was assayed in the presence of D-Leu, we also observed the LD-TPase products from exchange reactions, i.e. muropeptides with D-Leu at position 4 instead of D-Ala (muropeptides 13, 18, 21, 22, Figure 7B). Again, we observed muropeptides that arose from more than one Tlde1a activity. For example, muropeptide 12 has a 3-3 crosslink as well as a D-Leu modification, indicating that it has been generated by two different LD-TPase reactions.

We next quantified the different products of wild-type Tlde1a and Tlde1a variants with peptidoglycan. The results are summarized in Table S1 and are reported as molar amounts of the products that originated from LD-CPase activity, LD-TPase activity with a peptide acceptor, and LD-TPase activity with a D-Leu acceptor (Figure 7C). LD-CPase products prevailed over LD-TPase products for Tlde1a_WT_ and enzymatically active variants in the absence of D-Leu. As expected, Tlde1a_C113A_ was inactive and we also observed that the Tlde1a_G130Q_ and Tlde1a_W93A_ variants were inactive, while Tlde1a_R115A_ showed low activity. These results demonstrate the importance of the exchange pocket (Tlde1a_R115A_ and Tlde1a_G130Q_) and catalytic pocket (Tlde1a_W93A_) for all observed Tlde1a activities. Tlde1a_D40A_ produced less LD-CPase products than the wild-type enzyme, while the LD-TPase products were comparable or slightly higher than for the wild-type enzyme. Although many of the assayed Tlde1a variants appeared to abrogate all enzymatic activity, we found that the Tlde1a_H129A_ variant decreases LD-CPase activity more than LD-TPase activity. As shown in Figure 7C, Tlde1a_H129A_ produced only half of the LD-CPase products and a small amount of LD-TPase products with peptide acceptor as compared to Tlde1a_WT_. However, Tlde1a_H129A_ was fully active in LD-TPase exchange reactions with D-Leu as an acceptor. This suggests that the D-amino acid exchange reaction uses a different acceptor binding site than the LD-TPase reaction with a peptide acceptor and supports our structural observations of an exchange pocket and a catalytic pocket.

## DISCUSSION

Our structural and biochemical analysis of Tlde1a has revealed several molecular features that provide the enzyme with its various peptidoglycan modifying activities. We have shown that Tlde1a possesses an LD-TPase fold that has been adorned with structural modifications similar to those observed in the *E. coli* LD-TPase LdtD (YcbB), and in the *H. pylori* LD-CPase Csd6. These include a dynamic capping subdomain (LD-TPase) and extended active site cleft relative to the core enzyme fold (loops I and III, LD-CPase) (Fig. 1 and Fig. 2). Additionally, co-crystal structures with Tlde1a bound to D-amino acids reveals a binding pocket that is not specific to D-amino acid sidechains, which likely facilitates LD-TPase exchange activity (Fig. 3). Furthermore, binding of D-Met induces a conformational change in both a crystallization peptide artifact bound in the active site cleft of Tlde1a and in the Tlde1a capping subdomain (Fig. 3 and Fig. S2). This dynamic behavior suggests a role of the Tlde1a capping subdomain in enzymatic activity. Using the artifact His-tag peptide as a guide, ligand docking experiments provide predictions for how a peptidoglycan tetrapeptide stem might bind in the Tlde1a active site cleft. This prediction in combination with our co-crystal complexes demonstrates that the Tlde1a active site contains two D-amino acid binding pockets separated by the catalytic cysteine. These two pockets are an exchange pocket to bind a D-amino acid for LD-TPase exchange activity and LD-TPase with a peptide acceptor, and a catalytic pocket to position D-Ala^4^ of a peptide stem for LD-CPase activity (Fig. 3 and Fig. 4). Finally, we test our structural predictions by site-directed mutagenesis revealing that the LD-TPase and LD-CPase activities of Tlde1a are functionally linked at the molecular level (Fig. 5 and Fig. 7). Moreover, we observed that the LD-TPase and LD-CPase activities of Tlde1a can work in concert to produce peptidoglycan modifications, including peptide-stem crosslinks with an exchanged D-amino acid (Fig. 7 and Fig. S5).

The observed structural homology of Tlde1a with the LD-CPase domain of Csd6 (Fig. 2B) was of specific interest as Csd6 resembles a LD-TPase fold but only LD-CPase activity has been reported (55,60). When compared to LD-TPase domains in other structures, Csd6 was observed to have a unique active site structure. Many LD-TPases have a partially buried oxyanion hole (active site Cys and Tyr residues) that is surrounded on either side by two solvent exposed pockets. These pockets serve to bind donor and acceptor peptide stems for crosslinking. Overall, this creates a channel that extends through the enzyme placing the donor and acceptor sites on opposite surfaces of the enzyme. For example, this is clearly observed in the structure of the LD-TPase LdtD (YcbB) (54). In contrast, both Csd6 and our Tlde1a structure appear to lack opposite facing peptide binding sites, with the oxyanion hole facing only one side of the enzyme into a large active site cleft. This structural arrangement infers that Csd6 and Tlde1a cannot bind two peptide stems at once making them unable to create peptidoglycan crosslinks. However, our biochemical data demonstrates that unlike Csd6, Tlde1a can in fact form small amounts of 3-3 crosslinks (Figure 7). Given that the dynamic capping domain of Tlde1a is a molecular feature shared with the LD-TPase YcbB that can form crosslinks, we hypothesize that Tlde1a can alter its conformation to bind two peptide stems. Namely, our crystal structures have captured a closed conformation of Tlde1a involved in free D-amino acid exchange.

The substrate binding site of Csd6 was also observed to have additional loop extensions relative to standard LD-TPase folds that are hypothesized to contribute to LD-CPase activity (55). Similarly, Tlde1a has loop extensions near helix α2 and the 3_10_ helix (η1) that are homologous to loop I and loop III of Csd6 (Fig. 2C). Interestingly, these loops make up the binding site for free D-Ala and D-Met in Tlde1a and D-Ala in Csd6 (PDBid 4Y4V) (55). The authors speculate that this D-Ala and a second D-Ala bound near the active site cysteine of Csd6 may mimic how this enzyme binds its peptide stem substrate. Although Tlde1a has homologous loop extensions, helix α2 and the 3_10_ helix (η1) in all three of our structures adopt significantly different conformations relative to Csd6 (Fig. 2B). Additionally, the bound configurations of the D-Ala and D-Met in Tlde1a differ from the D-Ala in Csd6. Namely, the sidechains of the Tlde1a D-amino acids face away from active site and make no significant contacts, whereas the methyl sidechain of the modeled D-Ala in Csd6 faces towards the catalytic residues. Furthermore, the D-amino acids in Tlde1a have their backbone amine facing towards the active site for a potential reaction. Given the orientation of the bound D-Ala in Csd6, the authors conclude that D-Ala represents a bound product. However, considering our co-crystal structures, docking, and biochemical data we hypothesize that for Tlde1a, the loop regions of helix α2 and the 3_10_ helix (η1) comprise a binding pocket for an amino acid to be exchanged.

Comparison of Tlde1a with Csd6 provides a molecular explanation for the LD-CPase activity of Tlde1a and suggests that the D-amino acid binding pocket of Tlde1a is for substrate binding rather than product release. Yet together this analysis fails to reveal how Tlde1a also possess LD-TPase exchange activity. However, upon comparing Tlde1a to the LD-TPase YcbB, we observe that Tlde1a also has a capping subdomain (Fig. 2C). Previous work with YcbB has shown that its capping subdomain is required for LD-TPase activity in antibiotic resistance. Furthermore, at the molecular level the capping subdomain is thought to bind and stabilize substrates in the active site cleft for proper reaction with the active site cysteine (54). Given the observed His-tag peptide bound in the active site of Tlde1a, and that both the artifact and the capping subdomain change conformation upon binding of D-Met, we hypothesize that like YcbB the capping subdomain of Tlde1a stabilizes a peptide stem for transpeptidase exchange activity and potentially for crosslinking. Moreover, this conclusion is supported by our *E. coli* toxicity assays in which mutation of W90A abrogates activity (Fig. 5). Notably, W90 is buried within the core of the capping subdomain and is likely necessary for its proper fold and dynamics. Our structural and functional analysis demonstrates that Tlde1a combines the D-amino acid binding pocket of Csd6 with the capping subdomain feature of YcbB to produce its LD-TPase exchange activity.

As Tlde1a has molecular features that we hypothesize give the enzyme its dual activities, we probed these features by site-directed mutagenesis to support our structural conclusions. Interestingly, we observed that in our *E. coli* toxicity assays all protein variants except Tlde1a_D40A_ completely inhibited Tlde1a activity (Fig. 5). Additionally, our *in vitro* peptidoglycan assays showed that each Tlde1a variant except Tlde1a_D40A_ also significantly reduced enzymatic activity. Tlde1a_D40A_ slightly reduced LD-CPase activity but had no effect on LD-TPase crosslinking or exchange. This suggests that the D40 sidechain may only perform an accessory role in binding a peptide stem. Moreover, all inhibiting protein variants except Tlde1a_H129A_ had an equal effect in the reduction of both LD-CPase and LD-TPase D-amino acid exchange activity. Interestingly, Tlde1a_H129A_ (Fig 7 and Table S1) showed wild-type level D-amino acid exchange activity, but greatly reduced LD-TPase peptide crosslinking activity and partially reduced LD-CPase activity. Our docking predictions highlight H129 as a residue critical for orienting the D-Ala^4^ in a tetrapeptide stem for cleavage. Additionally, H129 is only part of the catalytic pocket and not part of the exchange pocket. Together this adds support to our structural conclusions that Tlde1a has both an exchange pocket and catalytic pocket important for its enzymatic functions. However, most point mutation variants inhibited Tlde1a toxicity and both peptidase enzymatic activities (Fig 5 and Fig 7). From this we conclude that the observed D-amino acid binding pockets and the capping subdomain work in concert contributing to both the LD-CPase and LD-TPase activities of Tlde1a.

Based on previous Tlde1a LD-CPase and LD-TPase biochemical assays, we did not expect to observe peptidoglycan crosslinking activity (38). The difference in products could arise from how the current and past peptidoglycan substrates were processed and purified. Namely, previous experiments used HPLC purified muropeptide substrates obtained after digestion with mutanolysin (38), whereas our assays used largely unprocessed peptidoglycan fragments. Interestingly, using peptidoglycan revealed that Tlde1a can not only create peptidoglycan crosslinks making it a true LD-TPase, but as shown in Figure 7 it can create a variety of differently modified products. These experimental results are also supported by the His-tag artifact bound in the active site of Tlde1a (Fig. 3A) and our tetrapeptide docking experiments (Fig. 4). Namely, the structural and docking data provide evidence that Tlde1a can likely accommodate the binding of different lengths and sequences of peptide fragments. Overall, this appears to suggest that Tlde1a can attack a prey cell peptidoglycan from various angles maximizing its toxic effects.

The observation that TPases can also exhibit CPase activity has also been reported for the other major class of peptidoglycan crosslinking enzymes, the DD-TPases (or penicillin-binding proteins, PBPs). Specifically, the bifunctional glycosyltransferase-DD-TPases PBP1A and PBP1B show substantial DD-CPase activity, i.e. the removal of D-Ala from position 5 of a pentapeptide, in particular when assayed at mildly acidic pH and upon activation with their cognate activators (61-64). It has been proposed that the CPase activity ‘rescues’ a TPase domain that is acylated with the donor peptide in a situation when there is no acceptor peptide available for the TPase reaction (61). In addition, PBP1A and PBP1B perform D-amino acid exchange reactions, replacing the D-Ala at position 5 by non-canonical D-amino acids such as HADA or NADA (64). While this is reminiscent for what we observed for Tlde1a at position 4 of the tetrapeptide, there is no data available about how the different acceptor substrates bind to the PBP DD-TPases.

Initial characterization of Tlde1a revealed that it is part of a large family of related enzymes. Specifically, Tlde1a is one of three subfamilies of Tlde1 enzymes which includes Tlde1a, Tlde1b, and Tlde1c (Fig. S6 and supplementary alignment file 1). Tlde1 enzymes are widely distributed in Gram-negative bacteria, however Tlde1a appears to be found primarily in *Salmonella* and *Bordetella* species (38). However, although related by possessing the LD-TPase motif (<u>H</u>XX14-17-(S/T)HG<u>C</u>h) the Tlde1 subfamilies show significant sequence variation. In particular, the disulfide bonds do not appear to be universal across the three Tlde1 subfamilies and importantly only Tlde1a displays an insert region relative to Tlde1b and Tlde1c for the capping subdomain (Fig. S6). In fact, even the Tlde1a insertion sequences appear to vary. Based on our structural and biochemical data with Tlde1a, these variations would likely lead to a more general TPase activity given the lack of a capping domain. In contrast, both R115 and G130 appear to be conserved indicating that at least the D-amino acid exchange activity may be shared by all Tlde1 orthologs. However, these sequence and thus molecular variations suggest that each Tlde1 enzyme has been structurally adorned in a unique way for a specific bacterial competition event. Overall, our findings provide an example of how a canonical LD-TPase fold has been structurally altered in the evolutionary arms race of T6SS effectors.

## EXPERIMENTAL PROCEDURES

### Protein expression and purification

Protein co-expression constructs for Tlde1a with Tldi1a (*Salmonella* Typhimurium 14028s) were synthesized and subcloned by Genscript in the vector pETDUET-1. Tldi1a with and without a sec signal was placed tag-less in multiple cloning site 1 (NcoI/HindIII) and full-length Tlde1a with a C-terminal 6His-tag was placed in multiple cloning site 2 (NdeI/XhoI). A construct containing Tlde1a in pET28b without the immunity protein Tldi1a created previously was also used in this study (38).

Tlde1a was overexpressed in *Escherichia coli* BL21(DE3)Gold cells using a Luria-Bertani (LB) broth culture medium. Cells were grown at 37°C until they achieved an optical density at 600 nm of 0.6. Protein expression was induced by 1 mM isopropyl-β-D-thiogalactopyranoside (IPTG), and cells were further incubated for 20 h at 20°C. Next, the cells were centrifuged at 4200 rpm for 25 min and resuspended in wash buffer (50 mM HEPES pH 7.5, 500 mM sodium chloride and 20 mM imidazole). The cells were then lysed using an Emulsiflex-C3 High Pressure Homogenizer (Avestin) and centrifuged at 17500 rpm for 30 min. The supernatant was run through a nickel ion affinity chromatography column that had been equilibrated with the wash buffer. The protein was eluted with elution buffer (50 mM HEPES pH 7.5, 500 mM sodium chloride and 500 mM imidazole). Consequently, the elution was concentrated to 2 mL and applied to a Superdex 75 column (GE Healthcare) for size exclusion chromatography. The column was equilibrated with gel filtration buffer (50 mM HEPES pH 7.2, 250 mM sodium chloride, and 1 mM beta-mercaptoethanol (β-ME). Purification of Tlde1a wild-type and Tlde1a variants for peptidoglycan modification assays excluded the reducing agent β-ME and used Tlde1a cloned within pET28b. To confirm high purity, all Tlde1a samples were run on a 12% SDS-PAGE gel and visualized by staining with Coomassie dye.

### Protein Crystallization

Crystallization of purified Tlde1a was screened for with commercially available screens (NeXtal), using a Crystal Gryphon robot (Art Robbins Instruments). Crystals were grown in 1:1 gel filtration buffer with 10 to 24 mg/mL Tlde1a, 0.2 M ammonium sulphate, and 20% PEG 3500 by sitting vapor-drop diffusion at 4 °C.

### Data collection and refinement

Tlde1a native, Tlde1a iodide derivative, and Tlde1a D-methionine datasets were collected in-house at 93K using a MicroMax-007 HF X-ray source and R-axis 4++ detector (Rigaku). The dataset for Tlde1a soaked with D-alanine was collected at the Canadian Light Source beamline CMCF-BM (08B1). Protein crystals were first cryo-protected with the addition of PEG 3350 or 6000 at a concentration of 35%, and then flash frozen in liquid nitrogen. The Tlde1a iodide derivative dataset included the addition of 500 mM NaI (Sigma) to the cryo-buffer and was incubated for 2 minutes. Co-crystal complex datasets with D-alanine (Sigma) and D-methionine (Sigma) included the amino acid added to the cryo-buffer at a concentration of 300 mM and 80 mM respectively, followed by an incubation time of 3 minutes. All the datasets were processed using XDS (65) and CCP4 (66). Phases for Tlde1a were obtained by single anomalous diffraction (SAD) using iodide as a heavy atom. Initial phases were calculated and the resulting protein model built using Phenix (67). Co-crystal complex structures were solved using Tlde1a (PDBid 7UMA) as a model for molecular replacement. All structures were further built using Coot (68) and refined using Phenix (67), Refmac5 (69), and TLS refinement (70). Molecular graphics were drawn with either UCSF Chimera (71) or ChimeraX (72).

### Tetrapeptide molecular docking

Docking of a peptidoglycan tetrapeptide stem with Tlde1a was performed using the server SwissDock (56,57). The native Tlde1a structure (7UMA) was used as the target structure after removing the CME adduct of residue C131 and the His-tag artifact. The selected peptidoglycan ligands (pubchem CIDs 25201603 and 134820170) were first prepared using the Grade Web Server (grade.globalphasing.org) to generate geometric restraints from the SMILES definitions. The pentapeptide CID 25201603 was modified to a tetrapeptide by removing the D-Ala^5^ using the program Avogadro (73). The tetrapeptides were not pre-docked by hand or other methods into the active site of Tlde1a before submission to SwissDock. Results were analyzed visually using UCSF Chimera and representative clusters chosen by SwissDock assigned rank, convergence of ligand conformations in each cluster, and alignment with the Tlde1a His-tag packing artifact.

### Construction of Tlde1 point mutation variants

Point mutations were created using the QuickChange II XL Site-Directed Mutagenesis Kit (Agilent Technologies) with either the pBRA SP-Tlde1_WT_ plasmid or Tlde1_WT_ pET28b plasmid used as template. Constructs were confirmed by sequencing.

### *E. coli* toxicity assays

Overnight cultures of *E. coli* DH5α carrying pBRA SP-Tlde1_WT_ or mutants were adjusted to OD_600nm_ = 1.2, serially diluted in LB (1:4) and 4 μL were spotted onto LB-agar (1.5%) containing either 0.2% D-glucose or 0.2% L-arabinose plus streptomycin and incubated at 37°C. Images were acquired after 20 h.

### Time-lapse microscopy of *E. coli* cells

For time-lapse microscopy, LB-agarose (1.5%) pads were prepared by cutting a rectangular piece out of a double-sided adhesive tape which was taped onto a microscopy slide as described previously (74). Overnight cultures of *E. coli* carrying pBRA SP-Tlde1_WT_ or mutants were sub-inoculated in LB (1:5) with 0.2% D-glucose and grown to OD_600nm_ = 0.6/0.7. Cultures were adjusted to OD_600nm_ = 0.35 and 1 μL of each strain was spotted onto LB-agarose pad containing 0.2% L-arabinose plus streptomycin. Images were acquired every 15 min at 30°C for 17 h using a Leica DMi-8 epifluorescent microscope fitted with a DFC365 FX camera (Leica) and Plan-Apochromat 63x oil objective (HC PL 614 APO 63x/1.4 Oil ph3 objective Leica). Images were analyzed using FIJI software(75). To determine cell length, approximately 30 cells were measured at the time point 4.5 h post-induction.

### Dynamic Light Scattering

Purified Tlde1a_wt_ and each variant was diluted in gel filtration buffer to 1 mg/mL. After dilution each protein sample was then filtered with a 0.1 μm cut-off membrane by centrifugation. Samples were loaded by capillary action into standard sensitivity capillaries and placed within a Prometheus Panta (Nanotemper). The instrument was allowed to stabilize at 20°C before measuring. Each Tlde1a protein sample was measured in triplicate, with 5 cumulants measured for each replicate. Scattering curves and apparent hydrodynamic radius were calculated using Prometheus Panta software with solvent viscosity estimated by the solute composition of the gel filtration buffer.

### Tlde1a enzymatic assays

In a final volume of 50 µl, purified Tlde1a proteins (10 µM) were incubated with peptidoglycan from *E. coli* BW25113Δ6LDT(76) in 20 mM HEPES/NaOH pH 7.5, 50 mM NaCl, with or without D-Leu (1 mM) for 4 h in a thermal shaker set at 37°C and 900 rpm. A control sample contained peptidoglycan and D-Leu but no protein. The reactions were stopped by heating at 100°C for 10 min. An equal volume of 80 mM sodium phosphate pH 4.8 was added and samples were incubated with 10 μg cellosyl (Hoechst, Frankfurt am Main, Germany) for 16 h in a thermal shaker set at 37°C and 900 rpm. Samples were boiled for 10 min and centrifuged at room temperature for 15 min at 16,000 × g. The supernatant was taken and the muropeptides present were reduced with sodium borohydride and as described (76). The reduced muropeptides were analysed by HPLC as published (59) but with a modified buffer, using a linear 140 min gradient from 100% 50 mM sodium phosphate pH 4.31 with 1 mg/L sodium azide (buffer A) to 100% 75 mM sodium phosphate pH 4.95, 30% methanol (buffer B). Eluted muropeptides were detected by their absorbance at 205 nm. The new muropeptides generated via Tlde1a reactions were collected and verified by LC-MS/MS as described (77). The results of MS analysis are shown in Supplemental Table S1. For the quantification in Figure 7C we calculated the relative molar percentage per PG subunit of the peptidoglycan modifications introduced by Tlde1a versions, i.e. the products from LD-CPase (CPase; free tripeptides), amino acid exchange reaction (TPase D-Leu; muropeptides with D-Leu) and transpeptidation reactions with peptide acceptors (TPase PG; 3-3 crosslinks) as follows: % modification (CPase, TPase D-Leu or TPase PG) = modified monomers (%) + 1/2 × dimers with 1 modification (%) + 1/3 × trimers with 1 modification.

## DATA AVAILABILITY

The X-ray structures and diffraction data reported in this paper have been deposited in the Protein Data Bank under the accession codes 7UMA, 7UO3 and 7UO8.

## SUPPORTING INFORMATION

This article contains supporting information.

## ACKNOWLEDGEMENTS

We would like to thank the laboratory of Jörg Stetefeld at the University of Manitoba and Nanotemper for access to a Prometheus Panta for DLS measurements. Additionally, we thank beamline CMCF-BM at the Canadian Light Source, which is supported by the Canada Foundation for Innovation (CFI), the Natural Sciences and Engineering Research Council (NSERC), the National Research Council (NRC), the Canadian Institutes of Health Research (CIHR), the Government of Saskatchewan. We thank Dr. Daniela Vollmer for the preparation of peptidoglycan from *E. coli* BW25113Δ6LDT. We also thank Dr. Gianlucca Gonçalves Nicastro for Tlde1a family alignment scripts.

## FUNDING AND ADDITIONAL INFORMATION

This work was supported by the Natural Sciences and Engineering Research Council of Canada (NSERC) grants RGPIN-2018-04968 to G.P., and a Canadian Foundation for Innovation award (CFI) 37841 to G.P. Work was also funded by the UKRI Strategic Priorities Fund (https://www.ukri.org) EP/T002778/1 to W.V and a São Paulo Research Foundation grant 2017/02178-2 to E.B-S.

## ABBREVIATIONS

β-ME: β-Mercaptoethanol
CME S,S: (2-hydroxymethyl)thiocysteine
CPase: carboxypeptidase
DLS: Dynamic light scattering
GlcNAc: N-acetylglucosamine
Hcp: haemolysin coregulated protein
IPTG: Isopropyl β-d-1-thiogalactopyranoside
LB: Luria-Bertani
mDAP: mesodiaminopimelic acid
MurNAc: N-acetylmuramic acid
PBP: penicillin-binding protein
PMSF: phenylmethylsulfonyl fluoride
SAD: single anomalous diffraction
SEC: Size exclusion chromatography
SP: signal peptide
SPI: Salmonella pathogenicity island
T6SS: type VI secretion system
TPase: transpeptidase
VgrG: valine glycine repeat G

