## Supplemental_data for "Molecular characterization of the type VI secretion system effector Tlde1a reveals a structurally altered LD-transpeptidase fold"

List of supplementary information:

Figures S1 to S6 (included in this PDF)
Table S1 (included as a separate file)
Alignment S1 (included as a separate file)
Supplemental movies Video S1 to S10 (included as separate files)

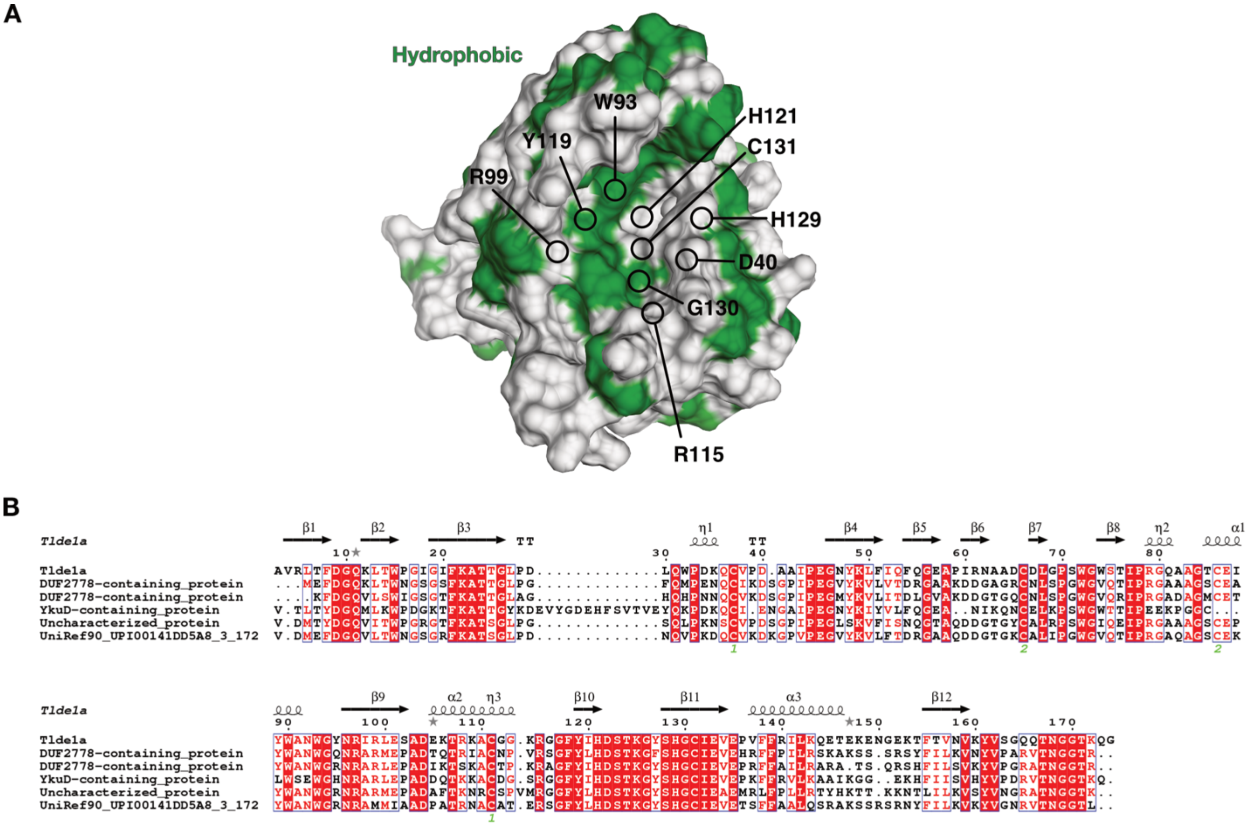


**Figure S1: Conservation of Tlde1a homologues.** A) Hydrophobic surface representation fo Tlde1a with conserved residues in the binding pocket indicated. B) Multisequence alignment of Tlde1a with the closest 5 relatives as calculated by the server Consurf (https://consurf.tau.ac.il/). Molecular graphics were drawn using UCSF ChimeraX (https://www.rbvi.ucsf.edu/chimerax/). Multisequence alignment was plotted by Espript (https://espript.ibcp.fr/).

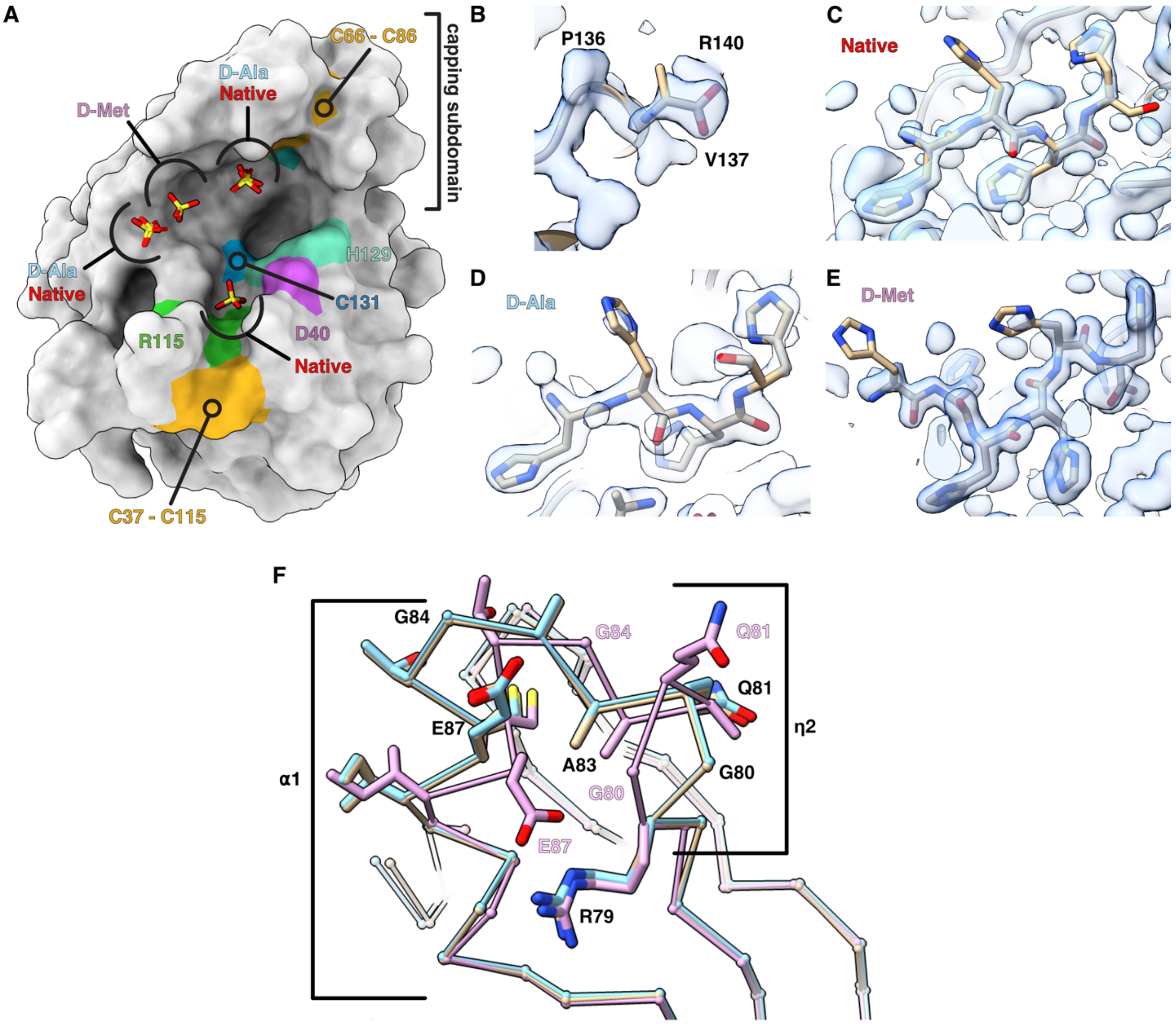


**Figure S2: Differences in ligand positions between Tlde1a crystal structures.** A) Surface representation of Tlde1a showing number of sulfates bound in the active site for each structure. The presence of D-Ala or D-Met alters the His-tag and sulfates. B) Second possible D-alanine binding site that occurs near a crystal packing interface in PDBid 7UO3. C) The observed electron density of the His-tag artifact when sulfate is bound (PDBid 7UMA), D) when D-Ala is bound (PDBid 7UO3) and E) when D-Met is bound (PDBid 7UO8). Each map is contoured at 1.2 rmsd. F) Tlde1a capping subdomain conformational changes. Bound to sulfate is shown in beige, bound to D-Ala in blue, and bound to D-Met shown in lavender. Observed conformational changes of the α1 helix and $3_{10}$ helix (η2) are highlighted. Molecular graphics were drawn using UCSF ChimeraX (https://www.rbvi.ucsf.edu/chimerax/).

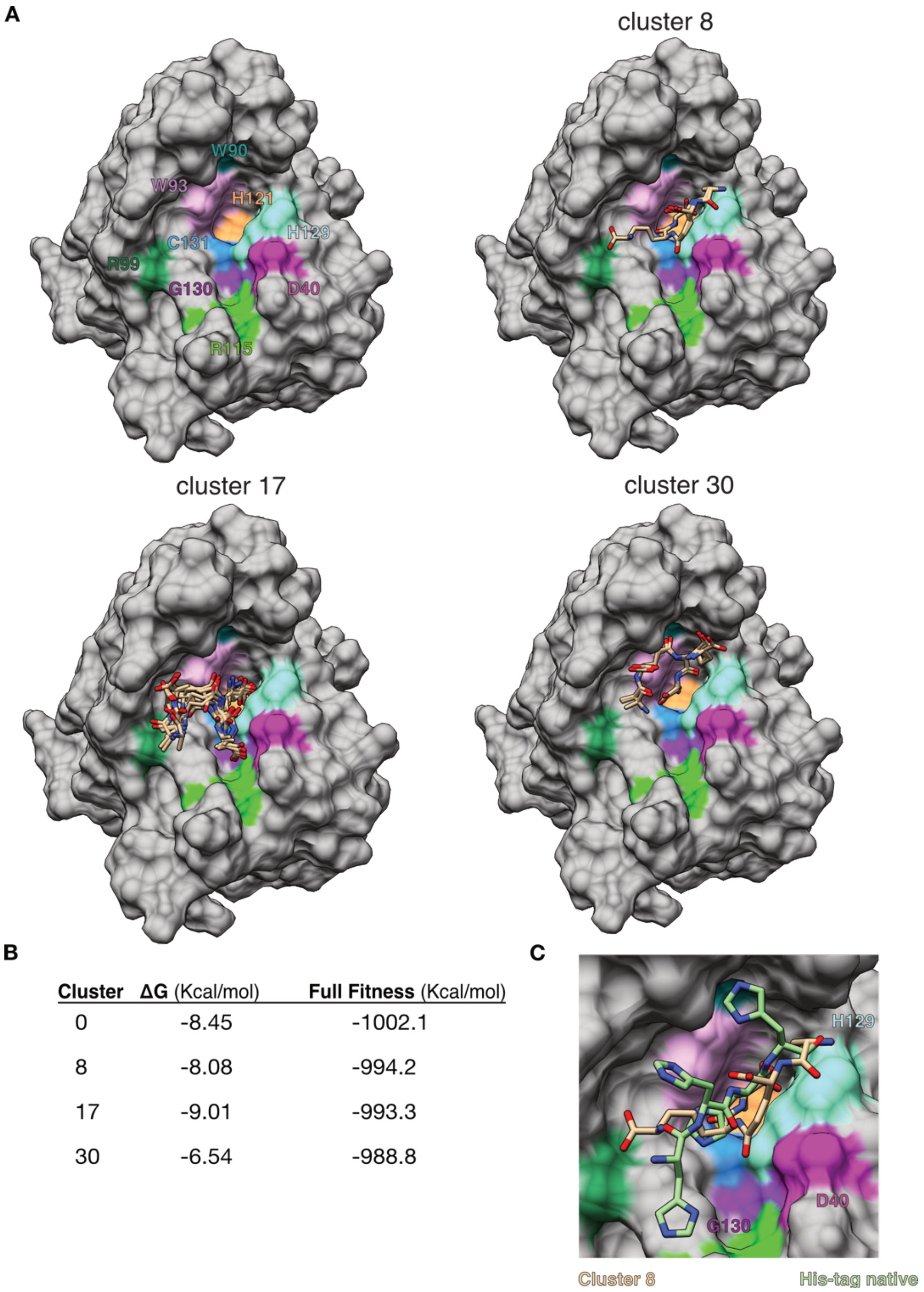


| Cluster | ΔG (Kcal/mol) | Full Fitness (Kcal/mol) |
|---|---|---|
| 0 | -8.45 | -1002.1 |
| 8 | -8.08 | -994.2 |
| 17 | -9.01 | -993.3 |
| 30 | -6.54 | -988.8 |

**Figure S3: Top three clusters chosen from molecular docking of the tetrapeptide L-Ala$^1$-γGlu$^2$-mDAP$^3$-D-Ala$^4$** A) Surface presentation of Tlde1a with conserved binding pocket residues highlighted by color. Each structure is labeled by cluster and shows the overlay of all cluster members with D-Ala indicated to show the tetrapeptide orientation. B) Docking statistics for each cluster as determined by SwissDock (http://www.swissdock.ch/). C) Comparison of cluster 8 with the conformation of the His-tag in 7UMA when only sulfate is bound.

A

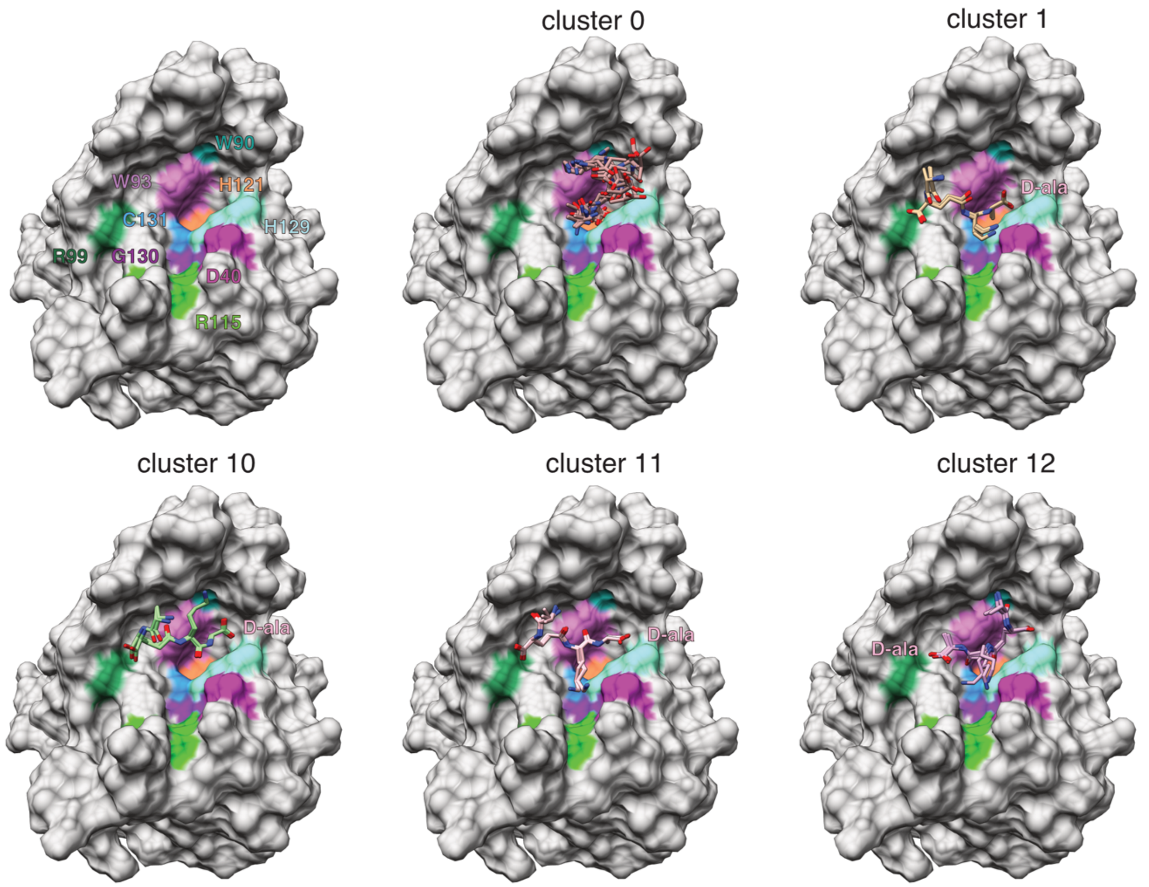


B

| Cluster | ΔG (Kcal/mol) | Full Fitness (Kcal/mol) |
|---|---|---|
| 0 | -8.66 | -1040.9 |
| 1 | -9.46 | -1038.1 |
| 10 | -10.32 | -1033.7 |
| 11 | -9.38 | -1033.0 |
| 12 | -10.88 | -1032.9 |

C

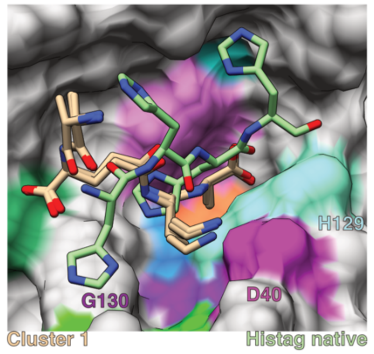


**Figure S4: Top five clusters chosen from molecular docking of the tetrapeptide L-Ala$^1$-γGlu$^2$-D-Lys$^3$-D-Ala$^4$** A) Surface presentation of Tlde1a with conserved binding pocket residues highlighted by color. Each structure is labeled by cluster and shows the overlay of all cluster members with D-Ala indicated to show the tetrapeptide orientation. B) Docking statistics for each cluster as determined by SwissDock (http://www.swissdock.ch/). C) Comparison of cluster 1 with the conformation of the His-tag in 7UMA when only sulfate is bound.

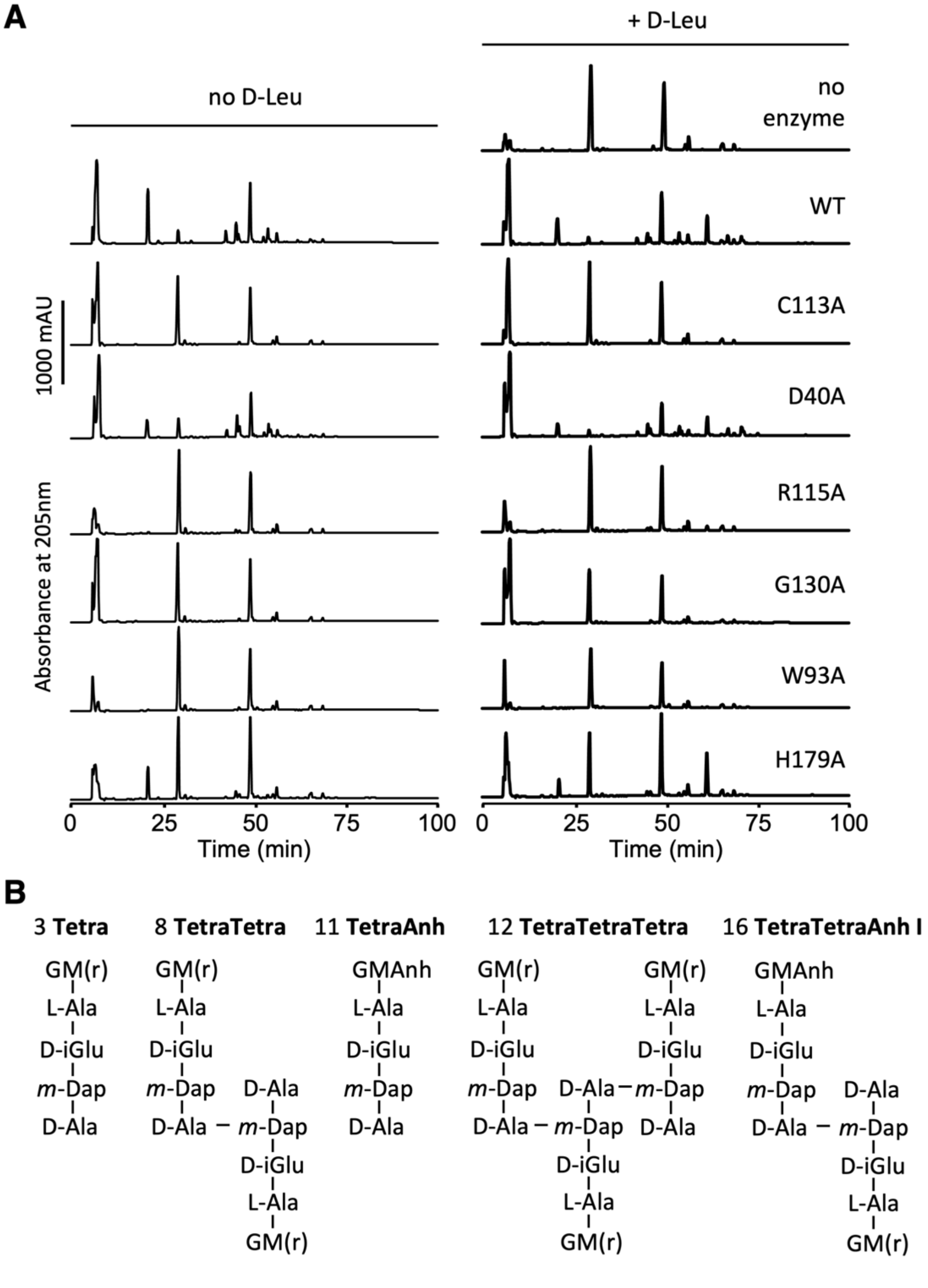


**Figure S5:** A) HPLC chromatograms showing the muropeptides released from reactions of Tlde1a variants with peptidoglycan from strain BW25113Δ6LDT in the presence or absence of D-Leu. A control reaction contained no enzyme and D-Leu. WT, wild-type Tlde1a. The different amino acid exchanges in Tlde1a versions are indicated on the right side. B). Structures of the main unmodified muropeptides, the numbers refer to peak numbers in Figure 7A. Figure 7B shows the structures of the modified muropeptide that were generated by Tlde1a.

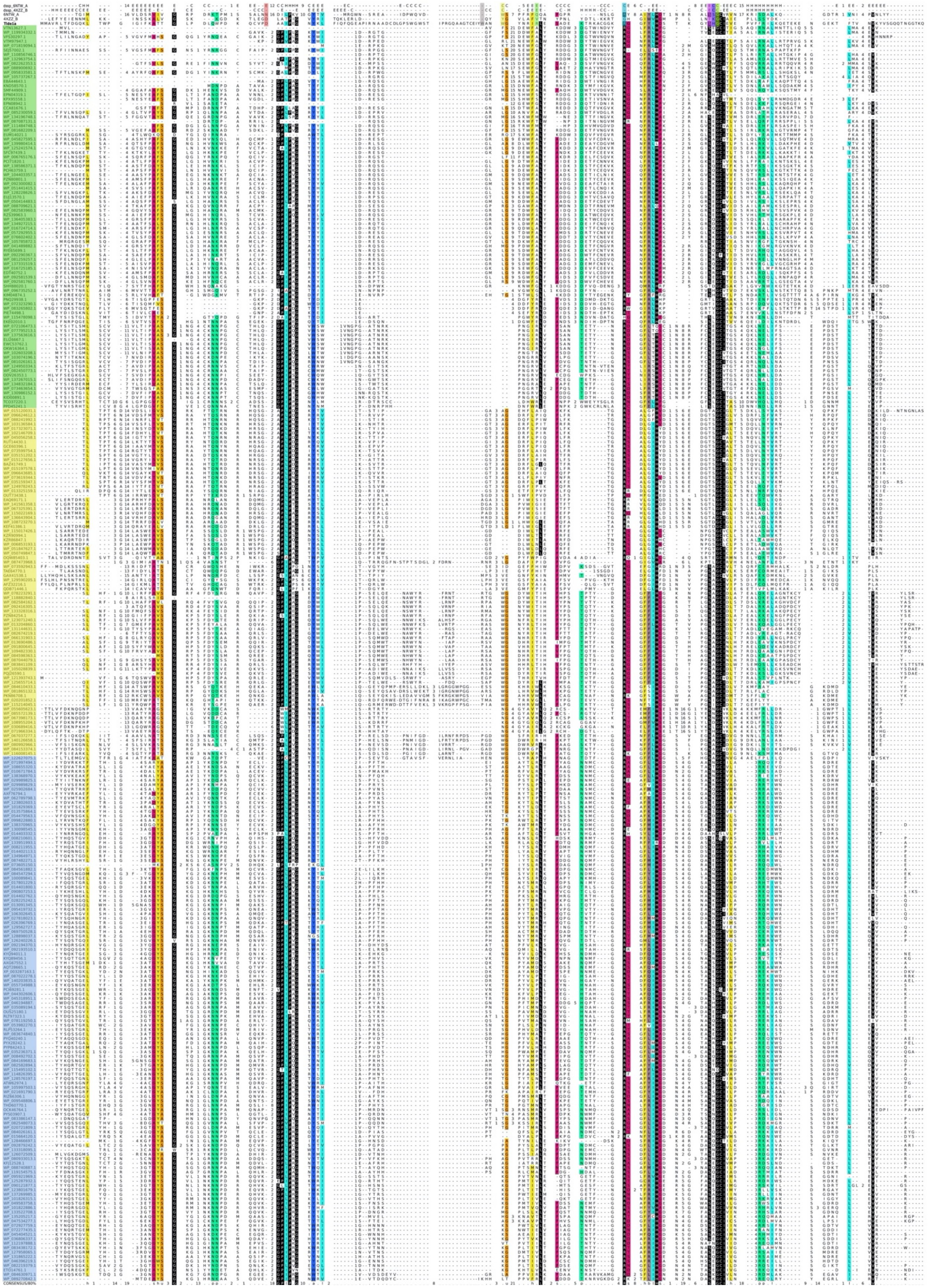

**Figure S6: Multiple sequence alignment of representatives from the Tlde1 family.** Alignment of representative sequences from Tlde1a (yellow), Tlde1b (blue), and Tlde1c (green) subfamilies retrieved from Sibinelli-Sousa et al. (2020). The secondary structure and sequences of *E. coli* L,D-Tpase YcbB (PDBid 6NTW) and *Helicobacter pylori* L,D-CPase Csd6 (PDBid 4XZZ) are shown at the top of the alignment. The consensus sequence is at the bottom of the alignment. Identical amino acid residues are colored in black and amino acids with similar properties are colored according to their side chains. Mutated Tlde1a residues shown in main Fig. 5 are indicated in the Tlde1a sequence at the top.

### Supplemental tables

**Table S1:** Quantification of muropeptides separated in Figure 7 and Figure S5. and results of the mass spectrometry analysis of selected muropeptides.

### Supplemental information files

**Tlde1 alignment S1:** Tlde1 sequence alignment file for Figure S6.

### Supplemental movies

**Video S1** – Time-lapse microscopy showing *E. coli* cells containing pBRA SP-Tlde1$_{WT}$ grown on LB-agarose (1.5%) pads with 0.2% L-arabinose (induced), related to Figure 5. Images were acquired every 15 min. Scale bar, 5 µm. Timestamps in hours:minutes.

**Video S2** – Time-lapse microscopy showing *E. coli* cells containing pBRA SP-Tlde1$_{C131A}$ grown on LB-agarose (1.5%) pads with 0.2% L-arabinose (induced), related to Figure 5. Images were acquired every 15 min. Scale bar, 5 µm. Timestamps in hours:minutes.

**Video S3** – Time-lapse microscopy showing *E. coli* cells containing pBRA SP-Tlde1$_{D40A}$ grown on LB-agarose (1.5%) pads with 0.2% L-arabinose (induced), related to Figure 5. Images were acquired every 15 min. Scale bar, 5 µm. Timestamps in hours:minutes.

**Video S4** – Time-lapse microscopy showing *E. coli* cells containing pBRA SP-Tlde1$_{W90A}$ grown on LB-agarose (1.5%) pads with 0.2% L-arabinose (induced), related to Figure 5. Images were acquired every 15 min. Scale bar, 5 µm. Timestamps in hours:minutes.

**Video S5** – Time-lapse microscopy showing *E. coli* cells containing pBRA SP-Tlde1$_{W93A}$ grown on LB-agarose (1.5%) pads with 0.2% L-arabinose (induced), related to Figure 5. Images were acquired every 15 min. Scale bar, 5 µm. Timestamps in hours:minutes.

**Video S6** – Time-lapse microscopy showing *E. coli* cells containing pBRA SP-Tlde1$_{R99A}$ grown on LB-agarose (1.5%) pads with 0.2% L-arabinose (induced), related to Figure 5. Images were acquired every 15 min. Scale bar, 5 µm. Timestamps in hours:minutes.

**Video S7** – Time-lapse microscopy showing *E. coli* cells containing pBRA SP-Tlde1$_{R115A}$

grown on LB-agarose (1.5%) pads with 0.2% L-arabinose (induced), related to Figure 5. Images were acquired every 15 min. Scale bar, 5 μm. Timestamps in hours:minutes.

**Video S8** – Time-lapse microscopy showing *E. coli* cells containing pBRA SP-Tlde1$_{H129A}$ grown on LB-agarose (1.5%) pads with 0.2% L-arabinose (induced), related to Figure 5. Images were acquired every 15 min. Scale bar, 5 μm. Timestamps in hours:minutes.

**Video S9** – Time-lapse microscopy showing *E. coli* cells containing pBRA SP-Tlde1$_{G130Q}$ grown on LB-agarose (1.5%) pads with 0.2% L-arabinose (induced), related to Figure 5. Images were acquired every 15 min. Scale bar, 5 μm. Timestamps in hours:minutes.

**Video S10** – Time-lapse microscopy showing *E. coli* cells containing pBRA SP-Tlde1$_{S128A}$ grown on LB-agarose (1.5%) pads with 0.2% L-arabinose (induced), related to Figure 5. Images were acquired every 15 min. Scale bar, 5 μm. Timestamps in hours:minutes.
